# Order-Selective Cells Tile Temporal Space and Predict Order Memory in Humans

**DOI:** 10.1101/2025.10.24.683575

**Authors:** Jie Zheng, Elisa C. Pavarino, Ivan Skelin, Mar Yebra, Jake Schwencke, Wenying Zhu, Chrystal M. Reed, Suneil K. Kalia, Taufik A. Valiante, Steven G. Ojemann, Daniel R. Kramer, John A. Thompson, Adam N. Mamelak, Gabriel Kreiman, Ueli Rutishauser

## Abstract

Remembering the temporal order of events is critical for episodic memory, allowing us to link individual events into sequences. While the medial temporal lobe and prefrontal cortex are essential for this process, the underlying neural mechanisms remain poorly understood. Here we characterized the representation of order information at the level of single neurons and field potentials recorded from human neurosurgical patients watching naturalistic videos of everyday events and later recalling the order and content of the events depicted. We found order-selective cells (OSCs) in the human hippocampus, amygdala, and orbitofrontal cortex that responded selectively to specific event orders, independent of event content or absolute time. OSCs exhibited transient theta phase precession following their preferred order during both memory encoding and retrieval, the strength of which predicted participants’ order memory accuracy. During retrieval, OSC spike timing relative to theta varied with the relative position of their preferred event within the recalled event sequence, enabling selective retrieval of relevant events. These findings reveal a neural substrate for representing, encoding into and retrieving from memory absolute and relative ordinal relationships between discrete events. OSCs tile temporal space into discrete ordinal positions, thereby weaving episodic experiences into coherent temporal narratives.

## Introduction

Memories, like scattered puzzle pieces, reveal the full picture of our past only when arranged in order. Memory for the order of events is a fundamental aspect of cognition, allowing individuals to organize experiences, make predictions, and engage in complex decision-making^1^. The ability to recall the order in which events occurred underpins daily activities, from recalling the steps of a recipe to following a conversation to making inferences about causality. Disruptions in order memory can lead to profound cognitive impairments, as seen in conditions such as Alzheimer’s disease and amnesia^2^, underscoring the importance of understanding the neural mechanisms that support this function.

Prior studies implicate the prefrontal cortex and medial temporal lobe in encoding and retrieving temporal information. The prefrontal cortex supports executive functions such as attention^3^, working memory^4^, and temporal organization^5^, which are essential for processing sequential information. Lesion studies^6^ show that prefrontal cortex damage impairs the ability to maintain temporal information in short-term memory. Likewise, the medial temporal lobe (MTL), particularly the hippocampus, is central to encoding and retrieving episodic memories, including their temporal order^7,8^. Neurophysiological recordings have revealed cells in the rodent and macaque MTL that encode aspects of time, including time cells^9,10^ that encode elapsed time since specific events and ramping cells in the entorhinal cortex and hippocampus that encode time over minutes^11,12^ or seconds^13,14^. However, it remains unclear whether these responses signaling time are related to order memory, or whether they alternatively are signaling time for other purposes. A major limitation of this body of work is that it focuses on the encoding of metric time, representing the continuous passages of time from seconds to minutes. But for many aspects of cognition, what is most relevant is the temporal relationship between discrete episodes, regardless of how long they last. Highlighting this gap, current metric-time based models cannot explain how the brain orders discrete episodes into structured memories^15^. Here, we seek to fill this gap by identifying neural mechanisms for encoding and retrieving the sequential order of discrete events and linking these neural mechanisms to order-based memory.

We recorded single-neuron activity and local field potentials from depth electrodes in 20 patients as they watched videos showing multiple discrete events (e.g., watering plants, cleaning tables). The movies were designed to closely approximate real-life experience, thereby allowing us to study order memory in an ecologically relevant context. Following movie watching, we assessed participants’ memory of the sequential content and temporal structure behaviorally. We center our analyses at event boundaries – critical timepoints where one event ends, and another begins – that determine the temporal structure of the memories^16^. This paradigm allowed us to examine how the brain detects and encodes the temporal structure of continuous experience while directly linking neural activity to behavioral measures of order memory in humans. This work provides a unique window into the cellular-level dynamics of temporal organization, offering insights that extend beyond the reach of noninvasive methods and are not readily accessible in animal models.

Our analyses revealed a novel functional class of neurons, Order-Selective Cells (OSCs), in the hippocampus, amygdala, and orbitofrontal cortex. The activity of these cells encoded the ordinal position of events independent of content or absolute time duration. Moreover, the timing of spikes of OSC cells relative to ongoing theta oscillations predicted participants’ accuracy on order memory tasks, indicating that a phase-dependent neural code, rather than firing rate alone, supports the sequential structuring of episodic memory. These findings uncover a neural mechanism by which discrete events are linked into coherent temporal narratives. By moving beyond continuous and metric time-based representations to event-based representations, our results provide evidence for order-based coding in the human brain, advancing our understanding of how episodic memories preserve not only content, but also the sequence that gives them meaning.

## Results

### Task and behavior

We studied how the order of events in a sequence shapes the formation and retrieval of memories for naturalistic movie clips with no sounds. The task consisted of three parts: encoding, scene recognition, and time discrimination. During encoding (Figure 1A), participants watched 25 different and novel custom-made video clips. Each clip consisted of a sequence of 4 events that were either separated by event boundaries (which coincide with a visual cut, Figure 1B and 1C) or appeared as one continuous shot (17 clips contained boundaries, 8 contained none). The event boundaries occurred at different time points (inter boundary interval: 8.00s ± 4.05s, mean ± s.d. throughout the text), with a wide temporal distribution across clips (Figure S1). Starting with the clip onset as event order 0, we refer to the events (movie clip parts) after each sequential boundary as order 1, order 2, and order 3. Each video clip was shown only once, and the content varied from one video clip to another (Figure S2).

**Figure 1.**
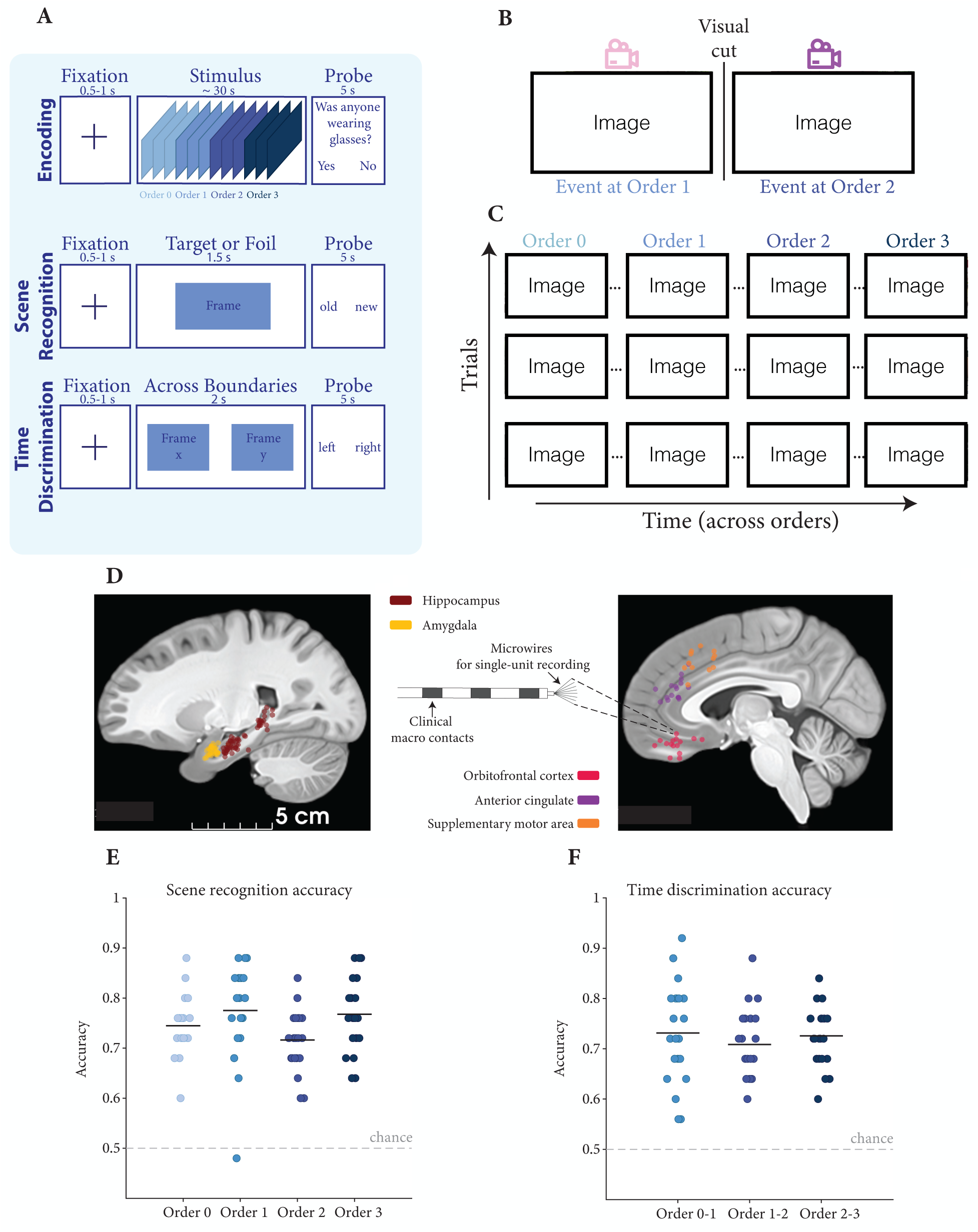
Experimental setup and behavioral performance. **A**, Experimental design. During encoding (top row), participants watched 25 unique videos (each played once, no sound) in a randomized order. Each video clip consisted of 4 events, referred to as event order 0, …, order 3 (denoted by different shades of blue, each lasting 7.34 ± 4.21 seconds, see also Figure S1), with a total duration of 30.09 ± 0.16 seconds. A two-alternative forced choice question was presented after each video clip to ensure participants’ engagement during encoding (e.g., was anyone wearing glasses in the clip?). After watching all the clips, patients performed two memory tests. First, during scene recognition (middle row), participants were shown one frame in each trial and were asked to indicate whether it was “old” (seen during encoding) or “new” (never seen before, the proportion of old was 50%) by pressing a key on the keyboard. Second, during time discrimination (bottom row), participants were presented with two previously seen frames in each trial and were asked to report which of the two frames occurred earlier (order memory) during encoding by pressing a key on the keyboard. The two frames were taken from the same movie clip but from different events. **B**, Example event boundary in one of the video clips. Here, the boundary is a visual cut due to a transition to a different point of view in the movie. **C**, Example frames after event boundaries at different ordinal positions (columns) for three different video clips (rows). Due to the bioRxiv’s policy, all the identifiable photographs in (B, C) have been replaced by the blank squares. These images are extracted from original videos produced by our team with media release consents and will be shown in the published version. **D**, Recording location of 97 microwire bundles (inset shows schematic of the electrode) across all 20 individuals. The hippocampus (crimson), amygdala (yellow), orbitofrontal cortex (red), anterior cingulate (purple), and superior frontal area (orange) are plotted on top of a sagittal view at x = 3.9mm of the brain template CIT168 (T1, 700um). Each dot represents the location of a microwire bundle with at least one neuron detected. **E, F**, Behavioral performance during the two memory recall experiments (E, scene recognition; F, time discrimination) across all 20 participants (each dot represents one participant). Responses are shown separately for each event order in E and for each order pair in F. Horizontal dashed line = chance level. Black solid lines = average performance. Almost all participants performed above chance levels.

Twenty patients with pharmacologically resistant epilepsy performed the task while we recorded the activity of single neurons (Figure 1D; Table S1 shows the patient demographics, and Table S2 shows the location of microwire bundles). One patient performed the task twice with a different set of stimuli for the second time, resulting in 21 data sessions in total. To assess participants’ engagement during encoding, a true or false question (e.g., was anyone wearing glasses in the clip?) appeared following every clip presentation. Participants answered 82 ± 11% of these questions correctly. After viewing all the clips, participants’ memory of events and their order in the presented clips was evaluated in two tests: scene recognition and time discrimination. During the scene recognition test (Figure 1A, middle), participants viewed a single static frame in each trial. These frames were chosen with equal probability from either the previously presented video clips (“targets”) or from other video clips that were not shown to the participants (“foils”). For each frame, participants made an “old” or “new” decision together with a confidence rating (1: sure, 2: less sure, and 3: very unsure). During the time discrimination test (Figure 1A, bottom), participants were shown two frames chosen from the same video clip they watched during encoding, presented side-by-side, and were asked to indicate which of the two frames appeared earlier in time together with a confidence rating (1:sure; 2: less sure; 3: very unsure).

In the scene recognition test, participants correctly identified 74.67 ± 6.73% of target images as “old” and 75.52 ± 7.01% of foil images as “new” (both above the chance level of 50%; Targets: *t*_20_ = 16.79, *p* = 3×10^-13^; Foils: *t*_20_ = 16.68, *p* = 3×10^-13^; one-sample *t*-test). In the time discrimination task, participants correctly identified which frame was shown first in 72.19 ± 4.18% of trials (above the chance level of 50%; *t*_20_ = 24.33, *p* = 2×10^-16^; one-sample *t*-test). Participants’ memory performance did not differ between frames following or across different event orders during time discrimination (Figure 1F, *F* (2, 40) = 0.454, *p* = 0.638, repeated ANOVA), while a moderate difference was observed in the scene recognition task (Figure 1E, *F* (3, 60) = 2.952, *p* =0.040, repeated ANOVA). The reaction times and confidence ratings were also similar for tested images across different orders during the scene recognition task (reaction time: *F* (3, 60) = 0.077, *p* = 0.972, repeated ANOVA; confidence rating: *F* (3, 60) = 0.207, *p* = 0.891, repeated ANOVA) and during the time discrimination task (reaction time: *F* (2, 40) = 0.807, *p* = 0.454, repeated ANOVA; confidence rating: *F* (2, 40) = 0.695, *p* = 0.505, repeated ANOVA). Overall, participants reliably remembered both the content of the clips and the relative temporal order of events, performing well above chance across all tasks and conditions.

### Neurons selectively responded to specific orders

We investigated the neuronal responses to event boundaries and their relationship to memory by recording single neuron activity from multiple brain regions, including the hippocampus, amygdala, orbitofrontal cortex, anterior cingulate, and supplementary motor area (the locations of microwire bundles with detected neurons are shown in Figure 1D). Across all areas, we recorded the activity of 965 neurons from 20 participants (see Figure S3 for spike sorting quality metrics). We first examined neuronal responses to event boundaries by comparing firing rates during a 0.5s-long post-boundary window to activity within the baseline period (i.e., the 0.5s window immediately preceding each boundary), pooled across all event boundaries regardless of serial position. For video clips with no boundaries, as a control, we aligned responses to virtual boundaries (i.e., same time point where event transitions are presented in other movies, see Methods), and compared responses between before and after these virtual boundaries. Figure 2A shows the response of an example neuron from the hippocampus that increased its firing rate around 300 milliseconds after event boundaries. No such change was observed in the clips without boundaries for the same example neuron (Figure S4A). Following our earlier work^17^, we refer to this type of neuron as a *boundary cell*. In all, 96/965 neurons (9.95%, *p* = 0.001, permutation test) were boundary cells. The regions with the largest proportion of boundary cells were the hippocampus and amygdala, with proportions significantly larger than expected by chance (hippocampus: 38/223, 17.04%, *p* = 0.001, permutation test; amygdala: 31/210, 14.76%, *p* = 0.001, permutation test). Boundary cells did not respond significantly to clip onsets (order 0; *p* > 0.05; Figure 2I, left).

**Figure 2.**
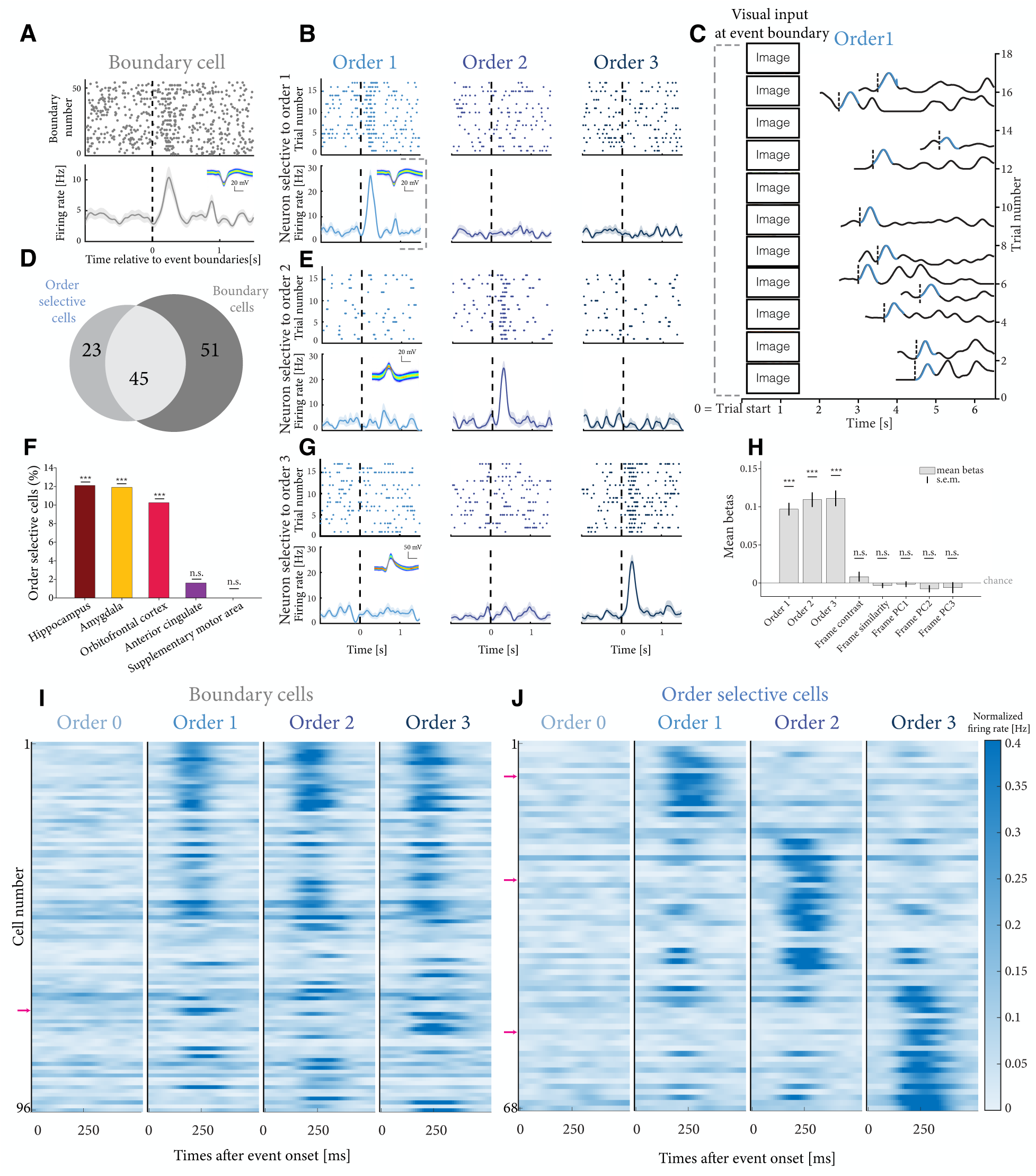
The activity of Order Selective Cells (OSCs) is preferentially tuned to a specific order of events and is invariant to content and absolute time. **A**, Example of a neuron that increased its activity after event boundaries. Each row of the raster plot (top row) is aligned to the event boundary (vertical dashed line, see examples in Figure 1B and 1C). Each line denotes a separate trial with a video clip consisting of 4 events, i.e., 3 event boundaries. Boundary cells are defined as neurons whose average firing rate significantly differed before boundaries ([-500, 0] ms) versus after boundaries ([0, 500] ms), considering all event boundaries regardless of their ordinal position (Wilcoxon rank-sum tests, *p* < 0.05). **B, E, G** Examples of three OSCs, responding preferentially to order 1 (B), order 2 (E), and order 3 (G). OSCs are defined as neurons whose average firing rate significantly differed before boundaries ([-500, 0] ms) versus after boundaries ([0, 500] ms) at a specific ordinal position (Wilcoxon rank-sum tests, *p* < 0.05), and also significantly differed across different orders (F statistic computed using the mean activities post event boundaries of the different orders, and compared to a shuffle test, *p* < 0.05; see Methods). The insets in A, E, G show the neurons’ waveforms obtained through the Osort spike sorting algorithm^38^ (see Methods). Note that (B) and (A) are the same neuron. **C**, Single-trial responses of the example neuron selective to order 1 (Figure 2B). Each row shows the firing rate of the neuron during viewing of a single video clip, aligned to the clip onset. In each row, the vertical dashed line indicates the time at which the first event boundary appeared, and the firing rate time course within [0, 500] ms following these event boundaries is highlighted in light blue. The interval [-500, 0] ms preceding the event boundaries is also shown for comparison. Trials for which the first event boundary happened beyond the limits on the x-axis are not shown. On the left in each trial, we show post-boundary frames. Due to the bioRxiv’s policy, all the identifiable photographs in (C) have been replaced by the blank squares. These images are extracted from original videos produced by our team with media release consents and will be shown in the published version. **D**, Number of OSCs, boundary cells, and neurons that are considered both boundary cells and OSCs. **F**, Percentage of OSCs in each brain area and its statistical significance against the chance level (see Methods). *** = *p* < 0.001, n.s. = not significant. **H**, Distribution of beta values from general linear models (GLM) describing each OSNs’ post-boundary firing rate using ordinal information and visual features as predictors (see Methods). The ordinal information was one-hot encoded (order 1, 2, or 3). The visual features include frame contrast, frame similarity, frame features from a computer vision model in the principal component space. Frame contrast is the standard deviation of each post-boundary frame in grayscale. The frame similarity is the cosine similarity score between the layer fc6 activation from AlexNet^18^ using inputs of pre-boundary frames and post-boundary frames. The principal components of post-boundary frames’ Alexnet fc6 representations were also computed. The bar heights represent the mean values of the beta coefficients averaged across all OSNs. The error bars represent the standard error of the mean. Chance level corresponds to a beta value of zero, indicated by the horizontal line. Statistical significance was computed by testing the betas for each predictor against the null hypothesis (Wilcoxon rank-sum test, *** = *p* < 0.001, n.s. = not significant). **I, J**, Summary plots of the neural activity of boundary cells (I) and OSCs (J) averaged across trials. Each row is a neuron, and each column is the average activity within [0, 500] ms after clip onsets (column 1) and order 1/2/3 boundaries (columns 2, 3, 4 respectively). The pink arrows indicate the example boundary cell shown in (A) and the 3 OSNs shown in (B, E, G). The color of the heatmap indicates the normalized firing rate (see scale on right).

Next, we asked whether neuronal responses varied with the serial order. We compared the responses to the three event boundaries, corresponding to the onset of the order 1, 2, or 3 events (Figure 1C). Many neurons exhibited pronounced response specificity, increasing their firing only following the event boundaries of one particular order, while showing little response to others (see examples in Figure 2B, 2E, and 2G). For example, the hippocampal neuron in Figure 2B exhibited a robust increase in firing following the first event boundary – consistently across nearly every video clip-but *not* after the second or third boundaries. Two additional examples include a hippocampal neuron selective for order 2 (Figure 2E) and a hippocampal neuron selective for order 3 (Figure 2G). We refer to neurons of this type as *order selective cells* (OSCs) - cells that show a significant response to at least one event boundary (assessed with a Wilcoxon rank-sum test comparing the firing rates 0.5s before versus after individual event boundaries) and, crucially, exhibit differential responses across boundaries at different orders (determined using a *F*-statistic along with the permutation test, see Methods). Sixty-eight of 965 neurons (7.45%, *p* = 0.001, permutation test) were classified as OSCs. The regions with the largest proportion of OSCs were the hippocampus (27/223, 12.11%, *p* = 0.001, permutation test), amygdala (25/210, 11.90%, *p* = 0.001, permutation test), and orbitofrontal cortex (8/78, 10.26%, *p* = 0.001, permutation test), with all proportions exceeding chance levels (Figure 2F). In contrast, only 2/124 (1.61%, *p* = 0.55, permutation test) in the anterior cingulate, and none (0/100, 0%, *p* = 0.99, permutation test) in the supplementary motor area met OSC criteria, not exceeding chance expectation (Figure 2F). Thus, OSCs were predominantly localized in the hippocampus, amygdala, and orbitofrontal cortex, which will be the focus of all subsequent analyses. Of the 68 OSCs, 45 also qualified as boundary cells, showing increased firing to multiple event boundaries but with modulation specific to event orders (Figure 2D). Similar to boundary cells, OSCs did not significantly change their firing rates following video clip onset (see ‘Order 0’ in Figure 2J, left; *p* > 0.05, permutation test) and in the clips without boundaries (Figure S4; *p* > 0.05, permutation test).

The response of OSCs was invariant to movie content. This can be appreciated by inspecting the rasters plots: unlike conventional raster plots, where each line represents a repetition of an identical stimulus, in Figure 2B, 2E, and 2G each line corresponds to a unique video clip with distinct visual content (example frames at event boundary 1 shown in Figure 2C). The response is highly similar despite distinct content. Visual features were significantly more similar *within* video clips than *across* different video clips (Figure S2; within vs across comparison: order 1: *p* = 2×10^-9^; order 2: *p* = 3×10^-9^; order 3: *p* = 2×10^-9^; Wilcoxon rank-sum test). Moreover, OSCs responded selectively to event boundaries, regardless of the absolute time since clip onset (Figure S1). Figure 2C shows the same neural responses as in Figure 2B but aligned to trial onset rather than event boundaries, with increased neural activity highlighted in light blue and event boundaries indicated by vertical dashed lines. This alignment demonstrates that OSC activity is tied to event boundaries themselves, rather than to the mere passage of time.

To quantitatively assess whether selective responses to different event orders might be attributed to visual content, we extracted multiple visual features from frames at event boundaries, including contrast, frame-to-frame similarity, and the first 3 principal components of the features extracted from the penultimate layer (fc6) in a neural network trained for object recognition (AlexNet^18^, see Methods). Pre- and post-boundary visual features did not differ significantly among the event boundaries at different orders (Figure S2; *F* (2, 72) = 0.14, *p* = 0.86, one-way ANOVA). Using these visual feature measurements along with event order information as predictors, we constructed a generalized linear model to predict trial-wise firing rates of OSCs. Event order - but not visual features - significantly explained the firing rate variations of the OSCs (Figure 2H; order 1: mean beta = 0.097, *p* = 1.1×10^-12^; order 2: mean beta = 0.110, *p* = 9.9×10^-13^; order 3: mean beta = 0.111, *p* = 1.7×10^-12^, significances obtained through signed Wilcoxon tests against zero). These findings indicate that OSC activation is driven by the ordinal position of events within a sequence, independent of the event content or absolute timing.

### Order information can be decoded in individual trials

We investigated whether event order could be decoded from population-level neuronal activity across different brain regions: hippocampus (n = 223), amygdala (n = 210), orbitofrontal cortex (n = 78), anterior cingulate (n = 124), and supplementary motor area (n = 100). For each region, we constructed pseudopopulations from all recorded neurons across patients and trained a linear-kernel support vector machine (SVM) to decode ordinal position information (order 1, order 2, or order 3, chance = 1/3), using firing rates from the 0.5-second window following each event boundary. SVM performance was assessed using 5-fold cross-validation. Each SVM was trained on a matched number of features (neurons) per region, with trials randomly split into training (80%) and testing (20%) sets, and the procedure was repeated 100 times. Neurons within the hippocampus, amygdala, and orbitofrontal cortex showed significantly high decoding accuracy compared to the shuffled distribution (Figure 3A, solid back versus grey; hippocampus: *Z* = 12.19, *p* = 0.01; amygdala: *Z* = 12.09, *p* = 0.01; orbitofrontal cortex: *Z* = 12.18, *p* = 0.01; Wilcoxon rank-sum test). Event order decoding accuracy in these three brain regions exceeded chance levels in temporal proximity to event boundaries (hippocampus: Figure 3B; amygdala: Figures S5F; orbitofrontal cortex: Figure S5L). However, the decoding accuracy of event order in these brain regions dropped significantly after removing the OSCs (Figure 3A, solid black versus striped black; hippocampus: *Z* = 12.22, *p* = 0.01; amygdala: *Z* = 12.00, *p* = 0.01; orbitofrontal cortex: *Z* = 12.22, *p* = 0.01; Wilcoxon rank-sum test). The anterior cingulate showed low but statistically significant decoding accuracy over chance (*Z* = 2.93, *p* = 0.01; Wilcoxon rank-sum test), and its decoding accuracy did not differ after removing all the OSCs in this region (*Z* = 1.51, *p* = 0.07; Wilcoxon rank-sum test). The decoding accuracy in the supplementary motor area showed no significant departure from chance (*Z* = 0.36, *p* = 0.36; Wilcoxon rank-sum test) and, as expected, remained unchanged after removing OSCs (Z = 0.00, p = 0.50, Wilcoxon rank-sum test), given that there are no OSCs detected in this region.

**Figure 3.**
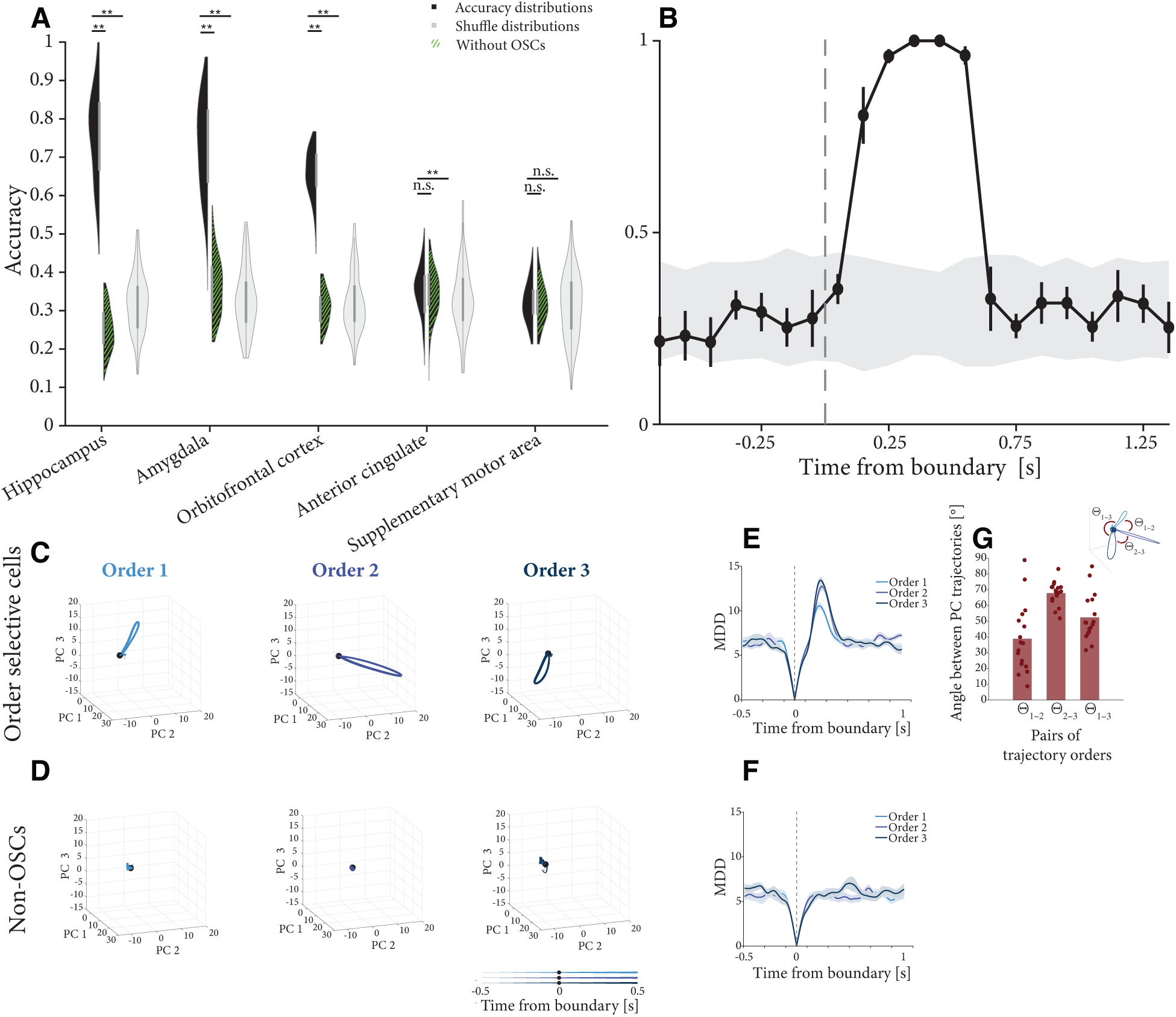
Order information is decodable in single trials from neural population activity. **A**, Accuracy of support vector machines trained to decode event order (1-3) from neuronal firing rates following event boundaries. Each SVM was trained using anatomically restricted pseudopopulations comprising all recorded neurons within a given brain region across patients. Features consisted of mean neuronal firing rates within the [0, 0.5]s window following event boundaries. The distribution of decoding accuracies across 5-fold cross-validation (repeated 100 times with different data splits) is shown in black violin plots. For comparison, parallel models were trained using identical features but with randomly shuffled order labels (gray violin plots), establishing chance-level performance. Additional analyses were performed using pseudopopulations that excluded OSCs within a given region (green striped pattern) to assess their contribution to order representation. Lower horizontal bars indicate statistical comparison between all neurons vs. neurons excluding OSCs; Upper horizontal bars indicate statistical comparison (see Methods) between all neurons versus shuffled controls and lower horizontal bars denote the statistical comparison between all neurons versus OSCs excluded. Asterisks report p-value thresholds, with * = *p* < 0.05, ** = *p* < 0.01. **B,** Decoding performance across time from event boundary trained on hippocampal OSCs. Each dot represents the accuracy of SVMs trained on 0.5-second windows, with a stride of 0.1 seconds. The accuracy of each time window is plotted at the middle time point of the time window. The decoding accuracy from the actual data is in black, and the decoding accuracy variability from the shuffled data is in grey. Error bars represent standard errors of the mean based on cross-validation. **C** and **D** Trial-averaged population trajectories in 3D principal component space, visualized across time ([-0.5, 0.5] seconds relative to event boundaries). These trajectories, derived through PCA of firing rates, represent the temporal evolution of hippocampal activity from either order-selective cells (C) or matched numbers of non-order-selective cells (D). Trajectories are plotted separately for each order (1, 2, and 3), with the black dot marking the time of event boundary occurrence. **E** and **F**, Multidimensional Euclidean distance (MDD) relative to event boundaries in the PC space (formed by all PCs that cover explained variance > 99 %) as a function of time aligned to the event boundary (i.e., time zero), computed using all OSCs (E) or non-OSNs in the hippocampus (F). The shaded area represents ± s.e.m. across trials. **G**, Quantification of angular separations between neural state trajectories in principal component space across different orders. For each trajectory, the main direction was computed through SVD decomposition, and the arccosine between two directions yielded their angle. The bar plot shows mean angles between order pairs, while scattered points represent angle measurements of individual trials.

We next examined how event orders are represented differently at the population level by evaluating the change in neural dynamics surrounding event boundaries using principal component analysis across groups of neurons of interests. Neural state representations underwent abrupt transitions immediately following boundaries for all three orders (Figure 3C; black dot indicates boundary time). These distinct neural state shifts were consistent with the changes in firing rates observed in boundary cells and OSCs reported in Figure 2. Similar neural state shifts were observed in the amygdala and orbitofrontal cortex (Figure S5). To quantify these state shifts, we computed the multidimensional Euclidean distance (MDD) in state space between each time point *t* and the boundary time. Plotting MDD as a function of time revealed an abrupt transition occurring within ∼300 ms post boundary (Figure 3E).

The neural state changes following different event orders shifted towards different directions (hippocampus: Figure 3H; amygdala: Figure S5C; orbitofrontal cortex: Figure S5I). To quantify the geometric properties of the trajectories, we measured the angular separation between trial trajectories generated by the different orders of events. For each order trajectory, we extracted the principal direction vector by performing singular value decomposition (SVD) and selecting the first component. Two complementary analyses both reveal order-specific consistency and inter-order distinctness. First, trajectories for each order traverse distinct regions of neural subspace, as measured by the non-zero angles between orders, seen for the hippocampus in Figure 3G, and for the amygdala and orbitofrontal cortex in Figure S5C and S5J, respectively. Second, when measuring angles relative to the first principal component axis, most orders showed non-random directions (order 1: 16.4° ± 49.1°, *t_16_* = 1.38, p = 0.19; order 2: 37.5° ± 24.6°, *t_16_* = 6.30, *p* < 0.001; order 3: 30.6° ± 8.6°, *t_16_* = 14.59, *p* < 0.001). This contrasts sharply with random directions (0.0° ± 60.5°). In contrast, when excluding OSCs from the analyses, the remaining neural population exhibited only small and essentially random changes as a function of time (hippocampus: Figure 3D, amygdala: Figure S5D, orbitofrontal cortex: Figure S5J), with MDD values showing no significant differences before versus after boundaries across event orders in the case of hippocampus and orbitofrontal cortex (hippocampus: *Z* =-0.33, *p* = 0.74; orbitofrontal cortex: *Z* = -0.33, *p* = 0.74, Wilcoxon signed-rank test), and mild but significant increase in the case of the amygdala ( *Z* = -2.93, *p* = 0.03, Wilcoxon signed-rank test). These analyses demonstrate that OSC trajectories of different orders consistently traverse highly structured, order-specific regions of neural state space.

### Theta power and theta phase precession are modulated by event orders

Phase precession of spiking relative to ongoing theta-band local field potential (LFP) is commonly regarded as a mechanism to encode temporal information. We next examined the properties of theta-band LFPs during video viewing. For each microelectrode, we computed theta power (2-10Hz) within a 1-second time window following each event boundary and preceding clip onsets (i.e., baseline). A microelectrode was classified as boundary-responsive if its averaged theta power following event boundaries differed significantly from baseline (*p* < 0.05; permutation test). Fifty-one microelectrodes met this criterion (25.8%, *p* = 0.008; permutation test), with 33 showing an increase (see example in Figure 4A) and 18 showing a decrease (see example in Figure 4C) in average theta power relative to baseline. Across all boundary-responsive microelectrodes, theta power following event boundaries increased (Figure 4B; *F* (3, 29) = 8.49, *p* = 3×10^-4^; one-way ANOVA) or decreased (Figure 4D; *F* (3, 14) = 12.11, *p* = 3×10^-4^; one-way ANOVA) progressively with the distance from the start of the video. As a control, we repeated the same analysis for the video clips with no boundaries, revealing no significant change in theta power as a function of (virtual) event order in the same microelectrodes (Figure S6B; *F* (3, 29) = 2.08, *p* = 0.125; Figure S6D: *F* (3, 14) = 3.09, *p* = 0.062; one-way ANOVA). Therefore, the change in power observed was not merely a function of time elapsed since video clip onset but rather due to the presence of event boundaries.

**Figure 4.**
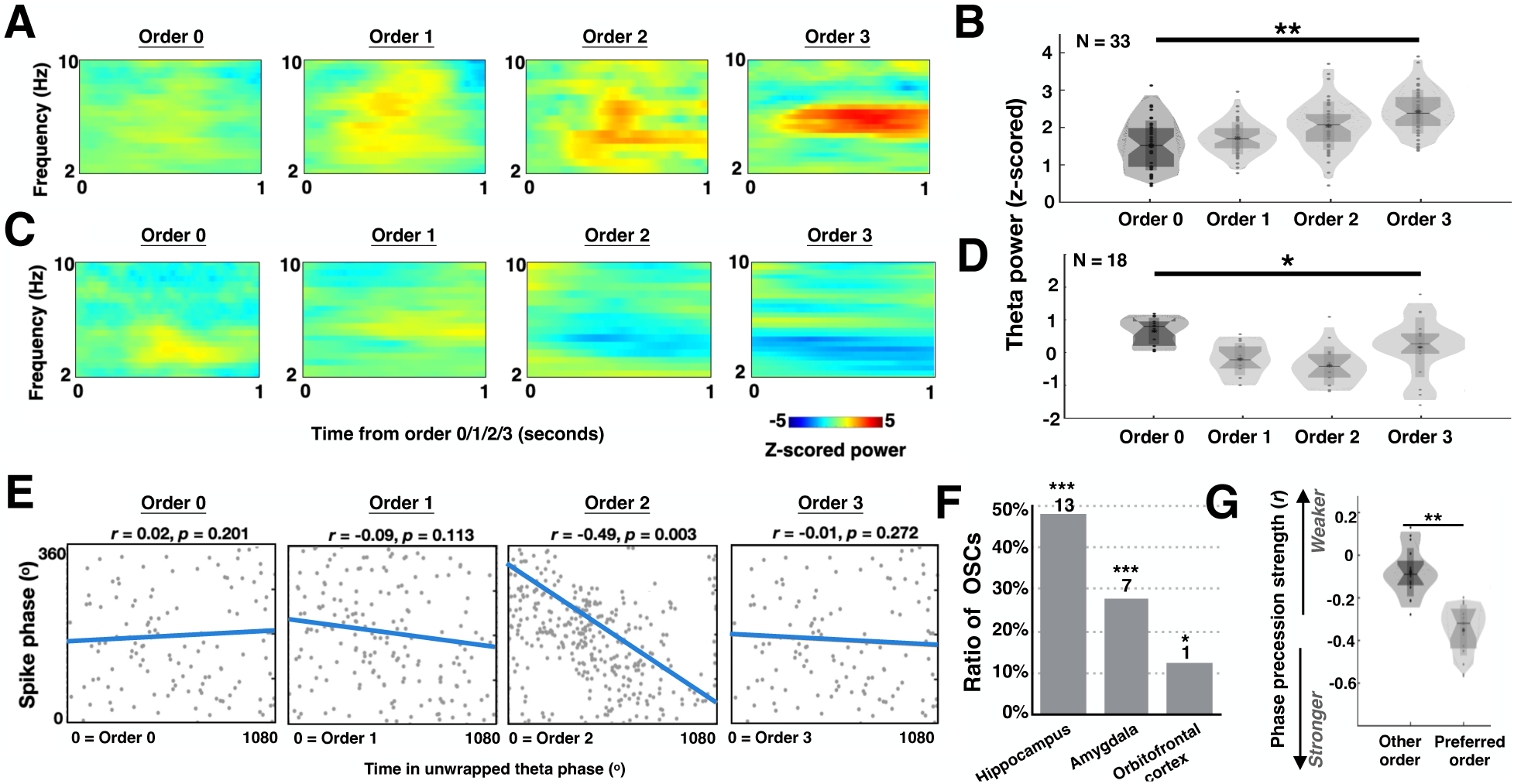
Theta power and phase precession are modulated by event order. **A** and **C**, Time frequency plots from 2 to 10 Hz for the local field potential signals from two example microelectrodes located in the hippocampus (**A**) and orbitofrontal cortex (**C**) aligned to the onset of event boundaries for order 0 to 3 (columns). Power within each frequency band is z-scored normalized to the baseline period (i.e., fixation cross in Figure 1A). **B** and **D**, Among microelectrodes that demonstrate significant theta power increase (**B**, n = 33 electrodes) or decrease (**D**, n = 18 electrodes) across different orders, the distribution of their normalized theta power computed within the 1-second time window following each event boundary for order 0 to 3. \**p* < 0.05, \*\**p* <0.01, ANOVA test. **E,** Example OSC located in the hippocampus showing theta phase precession following event boundaries at its preferred order (i.e., order 2). The y-axis shows the spike phase and the x-axis shows the unwrapped theta phase time (Methods); each dot denotes a spike. The strength of theta phase precession is quantified as the correlation coefficient (*r*) between spike phase and time in unwrapped theta phase. The more negative the correlation coefficients, the stronger the theta phase precession. **F,** Proportion of OSCs with significant theta phase precession following event boundaries for at least one order during encoding. \**p* < 0.05, \*\*\**p* <0.01, permutation test, see Methods. **G,** Phase precession strength of OSCs was strongest for the order for which they increased firing rate (“preferred”). \*\**p* <0.01, paired two-sided *t*-test.

On most boundary-responsive microelectrodes with theta power changes across event orders (38/51, 74.5%), we also recorded at least one OSC. We therefore evaluated whether the spiking activity of OSCs was modulated by theta-band LFP recorded from the same microelectrode. We started by assessing whether OSCs exhibit theta phase precession, a mechanism for sequence coding^19^. For each neuron, we first extracted the theta phase of the LFP from the same microelectrode at each spike time and aligned it to the elapsed time since the most recent event boundary. We quantified theta phase precession using the circular-linear correlation coefficient^20^ computed between the spike phase (circular variable) and elapsed time (linear variable^19,21–23^). To account for the non-stationarity of human theta ^23,24^, elapsed time was measured as the total accumulated theta phase after event boundaries rather than absolute time, a metric we refer to as ‘unwrapped theta phase’. Accordingly, 0° in the x-axis in Figure 4E corresponds to the boundary onset time, while 1080° marks the time after three theta cycles have elapsed. We then assessed the statistical significance of theta phase precession using a shuffle-based permutation test (*p* < 0.05, permutation test; see *Methods*) separately for each event order.

Twenty-one OSCs (21/68, 30.88%, above chance, *p* = 0.001, permutation test) spiked progressively earlier relative to ongoing theta (from 360° to 0°), characterized by a negative spike-phase correlation for at least one event order (see example in Figure 4E showing phase precession for event order 2). Most OSCs with theta phase precession following event boundaries were located in the hippocampus and amygdala (Figure 4F; hippocampus: 13/27, 48.15%, *p* = 0.001; Amygdala: 7/25, 28.00%, *p* = 0.002; permutation test). Across all 21 OSCs showing phase precession during encoding, the correlation coefficient was -0.31 ± 0.22 (*p* < 2 x 10^-8^, one-sample *t*-test), which is comparable to the strength of phase precession we reported previously following event boundaries^25^. Last, we asked whether the firing rate and phase precession response of OSCs were related. We did so by comparing phase precession strength between the event order to which a cell responded with a firing rate increase (“*preferred order*”) with the other event orders to which the cell did not change its firing rate (“*other order*”). Theta phase precession was stronger (i.e., more negative) following the OSC’s preferred order compared to other orders (Figure 4G; *t_21_* = 3.53, *p* = 1×10^-4^; two-tailed paired *t-*test). Therefore, most OSCs showed theta phase precession for the same event order for which they increased their firing rate, indicating a role for theta phase precession in order encoding.

### Theta phase precession during encoding tracks temporal relationships between events

Was the strength of theta phase precession during encoding related to the ability of participants to later recall the order of the events they had seen? During the time discrimination test (Figure 1A, bottom), subjects had to indicate the relative order of two previously seen frames. The two tested frames were always chosen from adjacent events (i.e., order 0 *vs* 1 events, order 1 *vs* 2 events, or order 2 *vs* 3 events). This design allowed us to test whether order retrieval accuracy was linked to the activity of OSCs during encoding. In other words, we asked whether OSCs’ activation at specific event orders during encoding predicted whether participants would later recall the correct temporal order of frames from those events during time discrimination.

The *preferred order* for each OSC was defined as the event order at which its firing rate was maximal. As shown in Figure 4G, most OSC cells also exhibited maximal theta phase precession at this same order. For each OSC, we categorized time discrimination trials into three conditions based on whether a frame from its preferred order event was present: (1) *preferred order leading trial* - frame from its preferred order event appeared earlier than the other tested frame in the original clip; (2) *preferred order following trial* - frame from its preferred order event appeared later than the other tested frame; (3) *other order trial* – neither tested frames came from the preferred order event. For instance, if an OSC’s preferred order was 1 (Figure 5A), then a *preferred order leading trial* would have tested frames extracted from order 1 and 2 events, a *preferred order following trial* from order 1 and order 0 events, and an *other order trial* from order 2 and order 3 events (Figure 5B-D). We then assessed theta phase precession strength during encoding separately for correct and incorrect responses to the order retrieval question across these three trial conditions. For example, Figure 5E shows an OSC with Order 1 as its preferred order, exhibiting a marked increase in firing rate and pronounced theta phase precession following event boundaries at this order. This cell showed stronger theta phase precession following order 1 event boundaries during encoding when the participant later retrieved the correct order in preferred order following trials (Figure 5E top), compared to those trials answered incorrectly (Figure 5E bottom). Across all 21 OSCs showing significant theta phase precession during encoding, the strength of theta phase precession was significantly stronger when participants correctly remembered the order of their preferred events either at the beginning (preferred order leading trails) or end (preferred order following trials) of the tested frames (Figure 5F left and middle; *t_21_* = 1.86, *p* = 0.037 and *t_21_* = 5.51, *p* = 1.5×10^-5^; paired two-tailed *t*-test). In contrast, for other order trials, theta phase precession strength was not predictive of retrieval accuracy (Figure 5F right: *t_21_* = 1.01, *p* = 0.16; paired two-tailed *t*-test). Further, the firing rate of OSCs during encoding by itself was not predictive of order accuracy (*t_21_* = 0.73, *p* = 0.24; paired two-tailed *t*-test).

**Figure 5.**
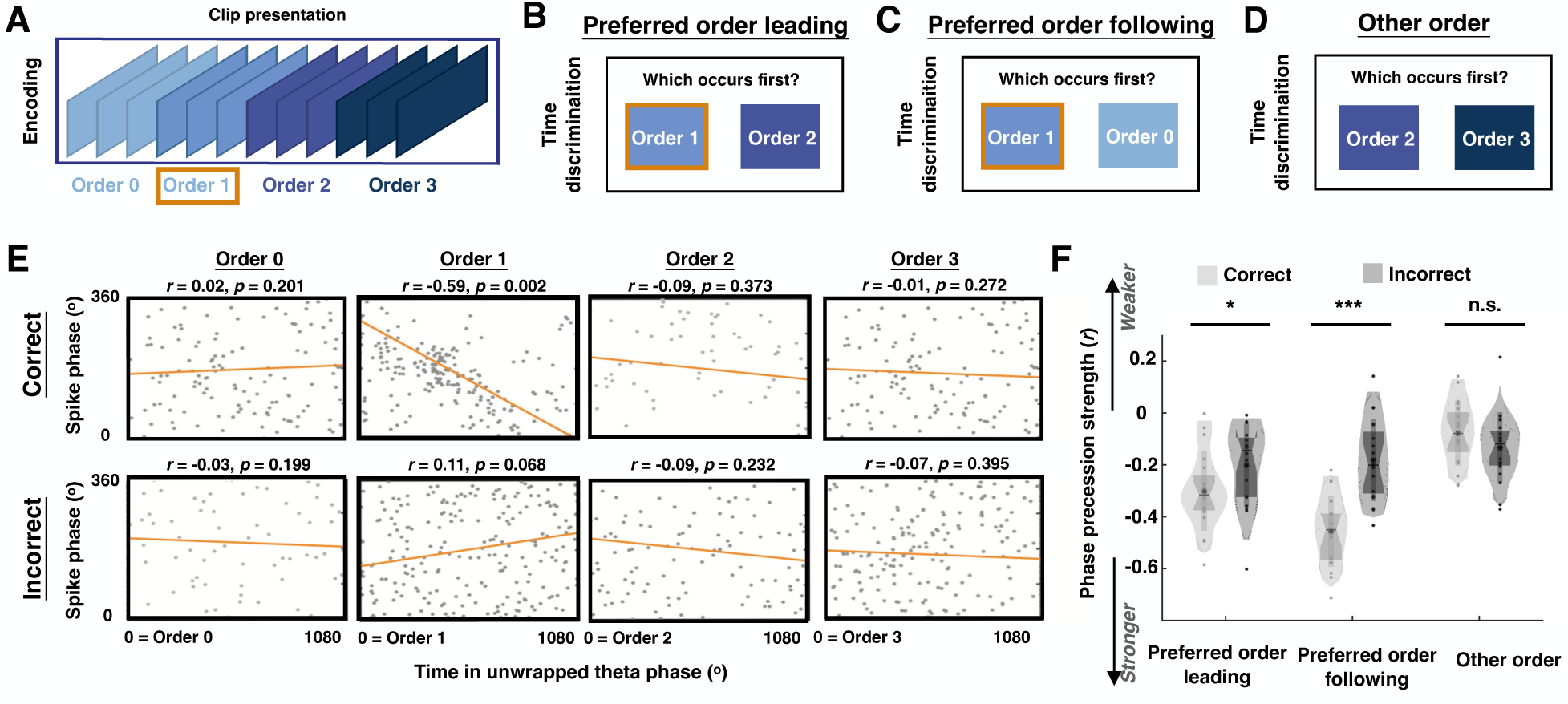
Theta phase precession during encoding predicts time discrimination accuracy. **A**-**D,** Trials during the time discrimination task (Figure 1A, bottom) are split into three conditions. As an example, we consider an OSC that fires more strongly during order 1 (preferred order). Time discrimination trials include frames with preferred order leading (**B**), preferred order following (**C**), or trials involving other orders (**D**). **E**, Theta phase precession during encoding of an order selective cell in the amygdala plotted separately based on the participant’s time discrimination accuracy (top: correct versus bottom: incorrect) for preferred order leading conditions during time discrimination (format as in Figure 4E). **F,** For OSCs that demonstrate theta phase precession during encoding (Figure 4F), this figure shows the distribution of theta phase precession strength following event boundaries at their preferred order, calculated based on participants’ time discrimination accuracy (correct versus incorrect), and plotted separately for preferred order leading, preferred order following and other order conditions.

Was theta phase precession also predictive of recognition memory accuracy? To address this question, we repeated the same analyses by categorizing encoding trials as ‘later remembered’ or ‘later forgotten’ (excluding trials with foil images). There was no significant difference in theta phase precession strength between remembered and forgotten stimuli drawn from an OSC’s preferred order events (*t_21_* = 0.58, *p* = 0.283; paired two-tailed *t*-test). This result also held for trials with tested frames extracted from OSCs’ non-preferred events (*t_21_* = 0.62, *p* = 0.270; paired two-tailed *t*-test).

### Theta phase precession during retrieval reflects temporal order memory

Our analysis so far has focused on phase precession during encoding. Do neurons also exhibit phase precession during order memory retrieval? To examine this, we aligned spike activity to image onset during the time discrimination task and computed the theta phase precession following trial onset using the same method as for encoding. We found that 24 of 68 OSCs (35.29%, *p* = 0.012, permutation test) showed theta phase precession during time discrimination, with an example cell shown in Figure 6A. This effect was mainly observed in OSCs recorded from the hippocampus and amygdala (Figure 6B; hippocampus: 12/27, 48.14%, *p* = 0.001; amygdala: 7/25, 28.00%, *p* = 0.007; permutation test). Across all OSCs, theta phase precession during time discrimination was significantly stronger if the trial included frames from OSCs’ preferred order events compared to when they did not (Figure 6C; preferred order leading *vs* other order: *t*_23_ = 4.37, *p* = 2×10^-4^, preferred order following *vs* other order: *t*_23_ = 5.62, *p* = 1×10^-5^, paired two-tailed *t*-test), indicating that phase precession during order memory retrieval was specifically related to order memory.

**Figure 6.**
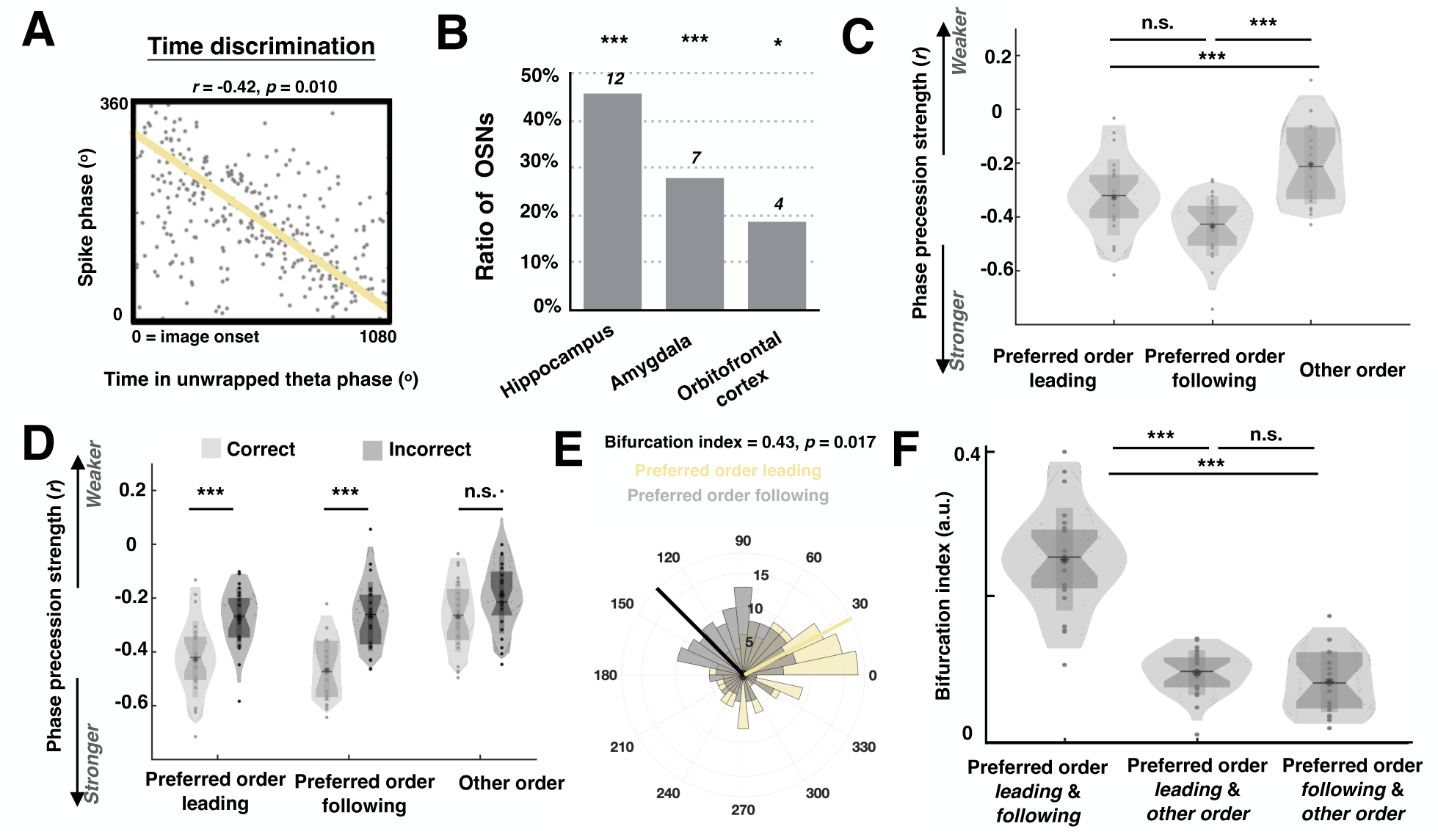
Spiking phases during memory retrieval reflect order memory outcomes. **A**, Example OSC in the hippocampus shows theta phase precession following image onset during time discrimination. **B**, Number and ratio of OSCs in the hippocampus, amygdala and orbitofrontal cortex showing significant theta phase precession following image onset during time discrimination encoding. \**p* < 0.05, \*\*\**p* <0.01, permutation test, see Methods. **C**, For all the OSCs showing theta phase precession during time discrimination in (B), theta phase precession strength is computed and plotted separately for preferred order leading, preferred order following, and other order conditions. **D**, For OSCs that demonstrate theta phase precession during encoding in (B), distribution of theta phase precession strength following image onset during time discrimination, calculated based on participants’ order memory outcomes (correct versus incorrect), and plotted separately for preferred order leading, preferred order following and other order conditions. **E**, Spiking phase (from image onset to button press) relative to theta rhythm of an example OSC in the hippocampus plotted separately when participants correctly recall the temporal order for preferred order leading (yellow) and preferred order following (gray) conditions. **F**, For all OSCs, bifurcation index computed using the spiking phase (from image onset to button press) relative to theta rhythm between preferred order leading and preferred order following conditions, or preferred order leading and other order conditions, or preferred order following and other order conditions.

We then asked whether the theta phase precession of OSCs during retrieval was related to retrieval performance, as we had observed during encoding. Within each condition (i.e., the preferred order leading, preferred order following, and other order condition, Figure 5B), retrieval trials were grouped according to whether participants correctly answered the order memory question or not. Theta phase precession following image onset was significantly stronger when participants correctly recalled the order of tested frames that included images from OSCs’ preferred order events (Figure 6D; Preferred order leading: *t_23_* = 5.24, *p* = 2.6×10^-5^; preferred order following: *t_23_* = 8.62, p = 7×10^-7^; paired two-tailed *t*-test). Consistent with the encoding results, theta phase precession strength did not predict retrieval accuracy for tested frames that did not contain images from OSCs’ preferred order events (Figure 6D; Other order: *t_23_* = 1.71, *p* = 0.101; paired two-tailed *t*-test). Further, the firing rate of OSCs during time discrimination alone was not predictive of order memory accuracy (*t_23_* = 0.83, *p* = 0.21; paired two-tailed *t*-test). Similarly, theta phase precession strength following image onsets during the scene recognition test showed no difference between trials when participants correctly recognized the target frames and those they forgot (*t_23_* = 0.44, *p* = 0.33; paired two-tailed *t*-test).

### OSCs encode relative ordinal position via spiking phase shifts

The time discrimination memory test evaluates participants’ memory for the relative temporal relationship between two adjacent events (e.g., did event A occur before event B?) rather than their absolute position within the sequence (e.g., was event A the second event in the sequence?). In contrast, OSCs encode absolute event order during encoding by increasing their firing rate (e.g., for the second event in the sequence; Figure 2E and 2J). This raises the question: how can OSCs that encode absolute event order during encoding support memory retrieval when participants are asked relative order questions in the time discrimination test? To address this, in a subset of time discrimination trials, the same image from an OSC’s preferred order event was paired with images that occurred either before it (preferred order following condition, Figure 5C) or after it (preferred order leading condition, Figure 5B). We asked whether OSCs encode the relative temporal relationships of their preferred order event differently for these two conditions. We first assessed the theta phase precession effect during time discrimination as reported in the previous section. When participants correctly answered the time discrimination questions, theta phase precession strength did not differ significantly from each other between the preferred order leading trials and preferred order following trials (*t_23_* = 1.36, *p* = 0.187, paired two-tailed *t*-test). This suggests that, while theta phase precession reflects the presence of the preferred order event, it does not by itself distinguish whether the event occurred earlier or later relative to its paired image, motivating a consideration of the following different theoretical model for retrieving the relative ordinal information during the time discrimination test.

Lisman and colleagues hypothesized that the sequential order of items can be represented and retrieved by neurons that are reactivated in sequence at distinct phases of theta-oscillations^26^. We reasoned that the preferred order event may occupy different ordinal positions when participants retrieve only a subset of the event sequence presented in the tested frames. For example, the order 1 event appears in the first position of the event 1-2 subsequence for preferred order leading trails (Figure 5B), while in the second position of the event 0-1 subsequence for preferred order following trials (Figure 5C). This raises the possibility that the spiking phases of OSCs encode different ordinal positions depending on the temporal subsequence being retrieved. To test this hypothesis, for each OSC, we computed the spiking phases relative to the theta rhythm across all the spikes from image onset to image offset during the order memory test. Note that we did not exclude periods of theta phase precession, which theoretically opposes theta phase coding/locking as theta phase precession is transient – occurring within 3 theta cycles or about 0.4 seconds after image onset – compared to the full analysis window of 2 seconds. Moreover, phase locking often emerges as a post-precession effect, as observed in place cells whose spikes stabilize at early theta phase near the end of phase precession, when the animal is about to complete traversal of the place field^21,27^. We therefore plotted the spiking phase distribution of the same OSCs for preferred order leading and preferred order following trials separately and computed the bifurcation index^28^ to quantify the differences in the phase that spikes locked to between these two trial types. The bifurcation index ranges from 0 to 1, with higher values indicating that the two distributions of phases had different means, and a value of 0 indicating that the phases are either not distinguishable or that neurons show no phase preference to begin with. For all the correct trials within the three conditions (preferred order leading, preferred order following, other order), we computed the bifurcation index for each pair of conditions. When participants correctly answered both the preferred order leading and preferred order following trials, OSCs tended to phase lock at different theta phases (example in Figure 6E; group analyses in Figure 6F). The bifurcation index for this comparison was significantly higher (bifurcation index = 0.28 ± 0.69) than for preferred order leading versus other order trials (*t_23_* = 6.22, *p* =1×10^-7^; paired two-tailed *t*-test), or preferred order following versus other order trials (*t_23_* = 5.97, *p* =9×10^-6^; paired two-tailed *t*-test). These findings suggest that the phase of OSC spiking shifts according to the event’s position within a retrieved subsequence, such that the absolute order of the preferred order event can be inferred directly from its theta phase.

## Discussion

Humans remember episodic events as discrete units, yet how these memories are integrated into a chronological continuum remains unknown. While prefrontal and medial temporal regions have been implicated in temporal encoding and recall, the specific neural mechanisms underlying this process remain poorly understood. We investigated the encoding of the order of sequential events by recording single-neuron activity and local field potentials in 20 drug-resistant epilepsy patients as they watched 25 video clips, each composed of a sequence of four everyday events of variable length. Participants’ memory was tested with scene recognition and temporal order discrimination tasks. We recorded 965 single units across patients and identified neurons in the hippocampus, amygdala, and orbitofrontal cortex that responded selectively to specific event orders, regardless of event content and the absolute time at which the events occurred (OSCs). Event order was decodable from the neural activity in these three brain regions, with OSCs being the major contributors. At the mesoscale, theta power changed along with event sequences, with OSCs exhibiting theta phase precession at their preferred event orders. Phase precession strength of OSCs during both encoding and time discrimination predicted participants’ order memory strength as assessed behaviorally, thereby linking the activity of OSC to memory. These findings reveal a neural mechanism for encoding event order in human episodic memory, demonstrating how event structure sculpts neural dynamics.

How do the cells we identified relate to known types of cells encoding aspects of time? Previous studies have described neurons involved in representing temporal information. For example, *time cells* in the rodent hippocampus^29^ fire at specific moments in time independently of spatial location, a finding that has also been confirmed in non-human primates^30^ and humans^10^. The OSCs described here share some features with time cells, including that both exhibit theta phase precession. Yet OSCs also exhibit important differences with time cells: whereas time cells track absolute elapsed time within a trial or behavioral interval, OSCs encode the *ordinal position* of discrete events independent of elapsed time. This feature of the activity of OSCs is most similar to event-specific rate-mapping neurons^31^ in rodents, which also respond selectively to event timing rather than absolute time and generalize across contexts (different mazes or, for OSCs, different clips). However, event-specific rate-mapping neurons fire when rewards are delivered following completion of specific laps, whereas OSCs fire naturally during viewing of complex sequences without explicit spatial or reward cues, suggesting that hippocampal neurons can track ordinal information in non-spatial contexts. Moreover, OSCs show stimulus-invariant responses modulated by theta-phase timing during both encoding and retrieval, while rate-mapping neurons are largely rate-coded, and their relationship to oscillatory activity remains unclear. In contrast to event-specific rate-mapping neurons, which primarily signal preferred events during encoding and have no clear role in retrieval or memory processes, OSCs are directly tied to memory. Finally, OSCs integrate both firing rate and theta-phase shifts to represent the relative ordinal position of preferred events within retrieved subsequences, whereas event-specific rate-mapping neurons mainly signal the occurrence of preferred order during encoding, with unclear roles in retrieval. Together, these findings suggest that OSCs extend the concept of temporal coding from absolute time to relative ordinal structure, providing a mechanism for encoding event order in complex sequences and supporting flexible retrieval.

What is the relationship between OSCs and the boundary cells that we reported before^17^? Both functional cell types share a key feature: they respond to event boundaries during ongoing movie clips with increased firing rates, but they do not respond to clip onsets (Figure 2G and 2H). This suggests that the activity of OSCs and boundary cells is not driven solely by visual transitions, since the screen changes at clip onset do not elicit similar responses. The new contribution here is that OSCs exhibit selective responses to specific boundaries - either the first, second, or third boundary within a clip - indicating that their firing reflects higher-order cognitive processing rather than boundary detection. We propose that OSCs may compute their order-selective responses by integrating inputs from boundary cells, which signal event occurrence, and from hypothetical ramping cells, which track elapsed time within the clip. As shown in Figure S7, the top row schematically illustrates boundary cells (see examples in Figure 2A and 2I) that increase their activity in response to all boundaries irrespective of their order. The middle row illustrates neurons signaling the passage of absolute time within each trial. These signals may be previously published time integrators^12^, or may be broader ramping signals, such as the increasing and decreasing theta power here reported (Figure 4 A-D). A product of these two types of responses leads to order selectivity (bottom row). In this way, the brain could maintain temporal information at multiple levels: ramping cells provide a continuous measure of time, boundary cells mark the occurrence of discrete events, and OSCs index the sequential order of these events. The identity and location of the putative ramping cells remain unknown in the human brain and will be the focus of future investigations (we did not find such cells in the areas we recorded from).

Most OSCs we identified were located in the hippocampus, amygdala, and orbitofrontal cortex (Figure 2E). This aligns with prior evidence that both the hippocampus and orbitofrontal cortex are critical for temporal coding. For instance, time cells^9^ and event-specific rate mapping neurons^31^ have been found in the hippocampus, which functions as a “temporal filter” essential for encoding time-based relationships and retrieving them to reconstruct event sequences^9^. Patients with hippocampal lesions struggle with tracking temporal durations^32^ and recalling events in a temporally organized sequence^7^. In the orbitofrontal cortex, local field potentials have been shown to track the evolving structure of temporal events, with distinct oscillatory patterns supporting the encoding and retrieval of time-dependent information^5^. Patients with orbitofrontal lesions exhibit marked deficits in tasks requiring the use of temporal information for decision-making and adaptive behavior^6^. We propose that OSCs in the hippocampus and orbitofrontal cortex may serve as a neural substrate for order memory in these regions. The role of OSCs in the amygdala (Figure 2E) is less clear, as there is limited literature linking this region to order memory. In our previous work^17^, we identified boundary cells in the amygdala, a finding replicated here. By analogy, OSCs in the amygdala may arise from integration of temporal and boundary signals, similar to the mechanism proposed for the hippocampus (Figure S7). Given the well-established interplay between emotion and time^33^, OSCs in the amygdala might contribute to this process. Finally, OSCs across the hippocampus, orbitofrontal cortex, and amygdala exhibited notable differences. In particular, theta-band phase precession was most prevalent in hippocampal OSCs during both memory encoding (Figure 4F) and retrieval (Figure 6B), highlighting a potential region-specific role in temporal sequence processing.

The spike timing of OSCs during encoding is indicative of whether subjects later correctly recall the order of events, linking OSCs and theta phase precession to temporal order memory. Stronger theta phase precession of the OSCs corresponded to better subsequent temporal order memory, which is specific to trials with preferred order events involved (Figure 5F). This is analogous to place cells, for which phase precession primarily occurs within the place field^21^, whereas here the “place field” is in temporal space - the preferred order that OSCs responded to, and such order-specific theta phase precession can occur during both encoding and retrieval. Crucially, the order memory effect depended on spiking timing/phases rather than firing rates of OSCs, consistent with prior findings^23^ and theoretical models proposing that theta-modulated spike timing underlies temporal coding^26^. Consistent with this model, OSCs not only signal the occurrence of their preferred order events but also adjust their spike timing according to the relative position of the event within a retrieved subsequence (Figure 6E and 6F). Because participants often retrieve only subsets of the event sequence, the same preferred order event can occupy different ordinal positions depending on the temporal context of retrieval. The bifurcation of spike phases across preferred order leading versus preferred order following trials demonstrates that OSCs dynamically encode relative ordinal positions through distinct theta phases. This phase-based coding mechanism provides a neural substrate for flexible temporal order memory, allowing hippocampal and associated circuits to reconstruct complex event sequences while preserving the relative event timing.

In sum, we identified OSCs that signal the relative temporal order of events, independent of content or absolute time. Predominantly in the hippocampus, orbitofrontal cortex, and amygdala, OSCs exhibit theta-phase precession linked to preferred event boundaries. The strength of phase precession predicted behaviorally assessed temporal order memory, thereby directly linking the activity of OSCs to memory. We propose that by dynamically encoding event position within sequences, OSCs provide a neural mechanism for integrating discrete episodic events into a coherent temporal narrative, offering new insight into human episodic memory formation.

## Methods

### Task

Twenty patients with drug-resistant epilepsy participated in a task whereby they watched video clips and reported their memory for events during those clips. The task consisted of three parts: encoding, scene recognition, and time discrimination (Figure 1A). During encoding, participants watched 25 novel and silent clips that either contained event boundaries or did not (Figure 1C). These clips were created by us as following. Each clip contained 4 to 5 events of daily activity (e.g., folding laundry, making coffee, etc.), with its original version (V0) filmed as a continuous shot from two different camera angles. A group of healthy human participants watched these original clips (V0) and annotated the event transitions. Based on their annotations, visual cuts were inserted either at the annotated event transition time (V1) or in the middle of events (V2), alternating between the two camera angles. Each time, eight V0 clips, nine V1 clips, and eight V2 clips without any content overlap were selected from the clip pools and presented during the encoding. We defined V1 and V2 clips as boundary clips, and V0 clips as no boundary clips. For both boundary clips and no boundary clips, the annotated event transition times marked the onset of events at different orders (e.g., order 1 event, order 2 event, order 3 event), with the onset of the order 0 event defined as the clip onset. Analyses focused on orders 0-3 due to limited trials for order 4. To assess participants’ attention, a yes/no question related to the content of the clip appeared randomly after every clip. After watching all the clips, participants were instructed to take a short break, roughly 5 minutes, before proceeding to the memory tests. Participants first performed the scene recognition test and were instructed to identify whether frames shown were from watched clips (answer: “old”) or from clips not shown (answer: “new”). To generate the testing frames for scene recognition, we first extracted frames from each clip, one randomly pulled out from each event within the clip. We then kept half of these extracted frames as target frames and replaced the other half with foil frames from different clips played by different actors/actresses or different clips played by the same actors/actresses. The total number of target and foil frames was counterbalanced across events at different ordinal positions. After the scene recognition test, we evaluated participants’ memory about the temporal structure of the clip using a time discrimination test. In each trial, two frames separated by event boundaries at different ordinal positions (e.g., one frame from order 2 event, one frame from order 3 event) were extracted from the same video clip and were presented side by side. Participants were instructed to indicate which of the two frames (that is, ‘left’ or ‘right’) appeared first (earlier in time) in the videos they watched during encoding.

### Participants

Twenty patients (12 females, age = 38 ± 13 years old, see participants’ demographics in Table S1) with refractory epilepsy volunteered for this study and provided their informed consent. Participants, who are implanted with electrodes for seizure monitoring, performed the task while they stayed at the hospital. The study protocol was approved by the Institutional Review Boards at the Toronto Western Hospital, Cedars-Sinai Medical Center, and University of Colorado Anschutz Medical Campus. The location of the implanted electrodes was solely determined by clinical needs. We targeted a total of 20 patients, with the sample sizes similar to those reported in previous publications^17,24^.

### Electrophysiological recordings

Broadband neural signals (0.1 – 8000Hz filtered) were recorded using Behnke-Fried electrodes^34^ (Ad-Tech Medical, Wisconsin, USA) at 32KHz using the ATLAS system (Neuralynx Inc., Montana, USA). We recorded bilaterally from the amygdala, hippocampus, and orbitofrontal cortex, and other regions. Electrode locations were determined by co-registering postoperative CT and preoperative MRI scans using Freesurfer’s mri_robust_register^35^. Each Behnke-Fried electrode shank had 8 microwires (or 1 microwire bundle) on the tip and was marked as one anatomical location in Figure 1D if at least one neuron was detected. To assess potential anatomical specificity across different participants, we aligned each participant’s preoperative MRI scan to the CIT168 template brain in MNI152 coordinates^36^ using a concatenation of an affine transformation followed by a symmetric image normalization (SyN) diffeomorphic transform^37^. The microwire bundles across all the participants are illustrated on the CIT168 template brain in Figure 1D. The MNI coordinates for microwire bundles with at least one OSC detected are listed in Table S2.

### Spike sorting and quality metrics of single units

The recorded signals were filtered offline in the 300-to 3000-Hz band with a zero-phase lag filter. Spike sorting was performed offline using a semi-automated template matching algorithm Osort^38^, with manual supervision applied to remove spurious neural clusters arising from artefacts. We identified 965 neurons in this dataset across all brain areas considered. The quality of our spike sorting results was evaluated using our standard set of spike sorting quality metrics^39^ for all putative single neurons (Figure S3).

### Cell classification

#### Classification of boundary cells

the neural activity of each recorded neuron was aligned to event boundaries in the clips. Event boundaries at different ordinal positions within the same clips were treated as independent trials. There were 3 event boundaries per clip, across 25 clips, with a total of 75 trials. The mean firing rate was computed within a 0.5-second time window before and after each event boundary and compared using a Wilcoxon rank-sum test. Cells showing a significant difference (p < 0.05) were classified as boundary cells.

#### Definition of Order Selective Cells (OSCs)

To identify OSCs, the neural activity of each recorded cell was aligned to event boundaries and separated into groups by event order. Within each order (i.e., the activity aligned to order 1, order 2, and order 3), a Wilcoxon rank-sum test was performed to compare the mean firing rates in the 0.5 seconds preceding and following the boundary (test 1). To assess differences across orders, we then computed the mean firing rate within the 0.5-second post-boundary time and calculated the F-statistic. We generated a null distribution by randomly reassigning post-boundary data to the three orders 10,000 times and recalculating the F-statistic for each shuffle. Neurons were considered significant if the observed F-statistic exceeded the 95th percentile of the shuffled distribution (p < 0.05, test 2). A neuron was classified as order selective (OSC) if it passed both criteria: (i) at least one order showed significant modulation in test 1, and (ii) post-boundary activity differed significantly across orders in test 2.

#### Chance level of boundary cells and OSCs

To evaluate the experiment-wide false discovery rate for boundary cells and OSCs (Figure 2F), we conducted permutation tests. For each neuron, the average firing rate within a 0.5-second window before or after event boundaries was randomly assigned to either the pre-boundary or post-boundary group across trials. We then applied the same detection method described earlier to identify boundary cells and calculate their proportion in the shuffled dataset. This randomization process was repeated 1,000 times per neuron, generating a null distribution of boundary cell counts. A similar approach was used to assess OSCs. Here, the average firing rate within the 0.5-second window following event boundaries was randomly reassigned to order positions 1, 2, or 3 across trials, producing a shuffled dataset and corresponding null distribution of OSC counts. To determine statistical significance, the observed number of boundary cells or OSCs in the actual data was compared against their respective null distributions derived from the shuffled trials.

### General Linear Model

A generalized linear model (GLM) was employed to identify the most informative predictors of OSCs’ firing rates. For each OSC, we fitted a trial-wise GLM using the form below:

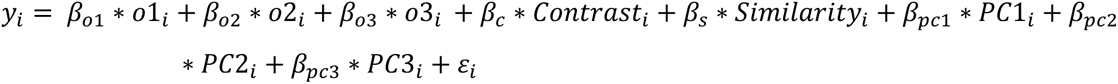

where *y_i_* denotes the firing rate of a neuron in the 0.5-second window following an event boundary on trial *i*. Predictors included the categorical order variables (*o*1, *o*2, and *o*3, one-hot encoded) and quantitative features of the video clips: stimulus contrast (calculated as the standard deviation of the grayscale video frame), frame-to-frame similarity (computed as the cosine similarity of AlexNet fc6 activation patterns of pre-boundary and post-boundary frames), and the first three principal components in AlexNet fc6 activation patterns of post-boundary frames. Continuous variables (e.g., firing rate, stimulus contrast, frame-to-frame similarity, etc.) were normalized by subtracting the mean and dividing by the difference between maximum and minimum values across trials. GLMs were trained using MATLAB’s glmfit.m function with a normal distribution and identity link, omitting the intercept term due to prior min–max normalization of the response variable. The resulting beta coefficients were averaged across OSCs (Figure 2H). Statistical significance of coefficients was evaluated with Wilcoxon signed-rank tests against the null hypothesis of median = 0.

### Classification analysis

#### Pseudopopulation feature matrix

To construct the feature matrix, we pooled anatomically restricted neurons across all participants. The matrix was organized such that each row represented an event boundary from a trial, and each column corresponded to a neuron from a specific anatomical region across participants. In Figure 3, we included only trials containing visual boundaries (i.e., V1 and V2 clips; see Task section under Methods), resulting in 17 trials. For each neuron within a given anatomical region, neural activity was aligned to event boundaries of three sequential events per trial. Each trial contributed three rows to the pseudopopulation matrix—one for each event order (1st, 2nd, and 3rd). The mean firing rate within a 0.5-second window following each boundary was computed and stored in the corresponding column for that neuron, with rows indexed by event order.

#### Decoding experiments

Support vector machines (SVMs) with a one-vs-all coding scheme were trained to decode event order based on firing rates averaged over 0.5-second windows following the boundaries of orders 1, 2, and 3. These analyses were conducted on anatomically constrained neural pseudopopulation feature matrices mentioned above (e.g., hippocampal neurons pooled across participants). To ensure consistency in feature count across brain regions, the smallest number of neurons available in any region was used to subsample features from each region. For each subsampled dataset, decoding accuracy was estimated using 5-fold cross-validation. This subsampling procedure was repeated 100 times, resulting in 100 decoding accuracy values per pseudopopulation. To establish a null distribution, the order labels were randomly shuffled for each subsampled dataset, breaking the link between firing rates and order identity. Decoding accuracy was then computed on these shuffled datasets using the same 5-fold validation, yielding 100 accuracy values for the null distribution. Statistical comparisons were performed using Wilcoxon rank-sum tests: (1) between the actual decoding accuracy distribution and the shuffled distribution, and (2) within pseudopopulations, comparing neurons with and without OSCs. Significance thresholds were set at \**p* < 0.05 \*\**p* < 0.01.

### Neural state space analyses

#### PCA analysis (average)

For each region, we constructed three neuron x time matrices aligned to order 1, 2, and 3 event boundaries. For each neuron, spike counts in a window around event boundaries ([-0.5, 0.5] s) were quantified into non-overlapping 10-ms time bins, smoothed using a Gaussian kernel (s = 200 ms), and averaged across trials. The three time series (one per order) were concatenated along the time axis, yielding a matrix where the columns (observations) correspond to the trials across 3 events, and the rows (features) correspond to the neurons. PCA was fit to the z-scored, transposed matrix, reducing the dimensionality of the neurons. The resulting principal component scores were then partitioned back to the three original event boundaries and analyzed both temporally and within the three-dimensional state space.

#### PCA analysis (trial-wise)

For single-trial analyses, we used the same preprocessing parameters ( [-0.5, 0.5] s windows around boundary events, 10 ms bins, Gaussian smoothing with s = 200 ms), without averaging across trials. For each region, three neuron × time matrices (each aligned to order 1, order 2, and order 3 boundaries) were computed by concatenating individual trials across the time axis, yielding a matrix where the rows correspond to neurons, and the columns represent concatenated time points across all trials and event orders. PCA was independently fit to this trial-wise, z-scored matrix. Multidimensional Distance (MDD): defined as the Euclidean distance between two time points in the principal component space. To estimate the standard error of the mean (s.e.m.), we calculated the MDD on a trial-by-trial basis and subsequently derived the mean and standard deviation of these individual trial measurements.

#### Neural trajectory angles

The principal direction for each trajectory was defined as the first right singular vector from an economical singular value decomposition (SVD). Angles between pairwise trajectories were computed as:

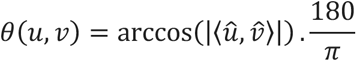

using normalized principal directions *û*, *v̂* and the absolute dot product to treat directions as undirected. To assess whether order-specific trajectories are significantly different from chance, we computed the directed angle between each trajectory direction and the x-axis. Further, we generated the null distribution by drawing 1000 independent vectors and measuring their angle with the x-axis. We report mean and standard deviation for real and null angles; significance was evaluated by a permutation-based p-value.

### Local field potential preprocessing

#### Pre-processing

Local field potentials (LFP) were recorded simultaneously with single neuron activity and were used for computing phase precession along with the spike activity from neurons detected from the same microwire. To eliminate potential influences of the spike waveform on the higher frequency parts of the LFP^40^, we replaced the LFP in a 3ms long time window centered on the detected spike by linear interpolation. We then downsampled this spike-free version of the LFP from 32kHz to 250Hz, followed by further post-processing using automatic artifact rejection^41^ and manual visual inspection by the first author of this paper using the function *fr_databrowser.m* in the Fieldtrip toolbox^42^ to remove trials with large transient signal changes from further analyses. Trials with large transient signal changes were removed from further analyses.

#### Spiking phases

We used a cycle-by-cycle analysis toolbox^43^ to detect the peaks, troughs and zero-crossings of theta waves. We then estimated theta phase at all points of time by linearly interpolating between the peaks (0° or 360°), troughs (180°) and zero-crossings (90° or 270°) cycle by cycle. Compared to a conventional Hilbert transform approach, this phase-interpolation method eliminates potential distortions introduced when estimating the phase of non-sinusoidal LFPs^44^. The phase assigned to a given spike was set equal to the phase estimated at the point at which the action potential was at its peak.

### Spiking phase-related analyses

#### Theta phase precession

We analyzed spikes that occurred within the first three theta cycles following boundaries during encoding or the onset of image display during time discrimination. Phase precession was quantified using circular statistics. For each neuron, we computed the circular-linear correlation coefficient between the spike phase (circular) and time in unwrapped theta phases (linear). Unwrapped theta phase was defined as the accumulated cycle-by-cycle theta phase starting at the alignment point (i.e., boundary onset or image onset). To assess statistical significance, we generated a null distribution for each circular-linear correlation using surrogate data generated from shuffling neurons’ spike timing 1000 times. This procedure maintained the firing rate and spike phase distribution of each neuron while scrambling the correspondence between the spike phase and spike time within each trial. A neuron was considered a significant phase precession neuron if the observed circular linear correlation exceeded the 95^th^ percentile (*p* < 0.05) of the surrogate null distribution of the correlation coefficient.

#### Chance level of neurons showing phase precession

We estimated the number of neurons exhibiting significant phase precession by chance by recomputing the spike-phase circular-linear correlation 1000 times using the surrogate data generated by shuffling trial numbers between the spike phases and LFP. For each iteration, we obtained the proportion of selected phase precession cells relative to the total number of neurons within each brain region. These 1000 values formed the empirically estimated null distribution for the proportion of phase precession cells expected by chance. A brain region was considered to have a significant amount of boundary cells or event cells if its actual fraction of significant cells exceeded 95% of the null distribution (Figure 4F and Figure 6B; *p* < 0.05).

#### Bifurcation index

For trials which participants correctly recall the order memory, the spike-phase distributions were constructed using all the spikes from the image onsets to button press during time discrimination and separately for each condition (e.g., preferred order leading versus preferred order following conditions). The separation between phase distributions was assessed using a circular distance metric and the Bifurcation index was computed as 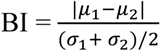, where *μ*_1_ and *μ*_2_ are the mean preferred phases for two conditions, and *σ*_1_ and *σ*_1_ are the corresponding circular standard deviations. A higher BI indicates stronger phase separation between conditions. Statistical significance of the BI was assessed using a permutation test, where condition labels were shuffled 1,000 times to generate a null distribution. A neuron was classified as exhibiting significant phase bifurcation if its observed BI exceeded the 95^th^ percentile of the null distribution.

### Statistical methods and software

Participants were not informed of the existence of event boundaries in the clips. All the statistical analyses were conducted in MATLAB, primarily using the Statistics and Machine Learning toolbox. For comparison against specific values, we used one-sample *t*-test. For comparison between two groups, we primarily used the Wilcoxon rank sum test, while for omnibus testing, we used one-way ANOVA, unless otherwise specified in the text. When the normality of data was not clear, non-parametric permutation tests were used to determine significance levels by comparing the real test-statistic to the null distribution estimated from the surrogate dataset.

## Data Availability

Data that support the findings of this study will be deposited at DANDI Archive upon acceptance.

## Code Availability

Codes that support the findings of this study will be deposited at GitHub upon acceptance.

## Acknowledgments

We thank the members of the Kreiman and Rutishauser laboratories for discussion and feedback, and all participants and their families for their participation. We thank Marielle L. Darwin for assistance with data acquisition, Jeffrey Chung, Lisa Bateman, the clinical teams at Cedars-Sinai Medical Center, Toronto Western Hospital, and University of Colorado Anschutz Medical Campus for patient care. This work was supported by National Institutes of Health grants U01NS117839 (to U.R.), K99NS126233 (to J.Z.) and R00NS126233 (to J.Z.). The funders had no role in study design, data collection and analysis, decision to publish or preparation of the manuscript.

## Author contributions

J.Z. conceived the project. J.Z., E.P., J.K., G.K., and U.R. contributed ideas for experiments and analysis. C.M.R, S.K.K., T.A.V., S.G.O., D.R.K., and A.N.M. managed the participants and surgeries. J.Z., I.S., M.Y., W.Z., J.A.T. collected the data. J.Z. and E.P. performed the data analysis. J.Z., E.P., G.K., and U.R. wrote the manuscript with input from all authors. U.R. and J.Z. acquired the funding.

---

**Figure S1.**
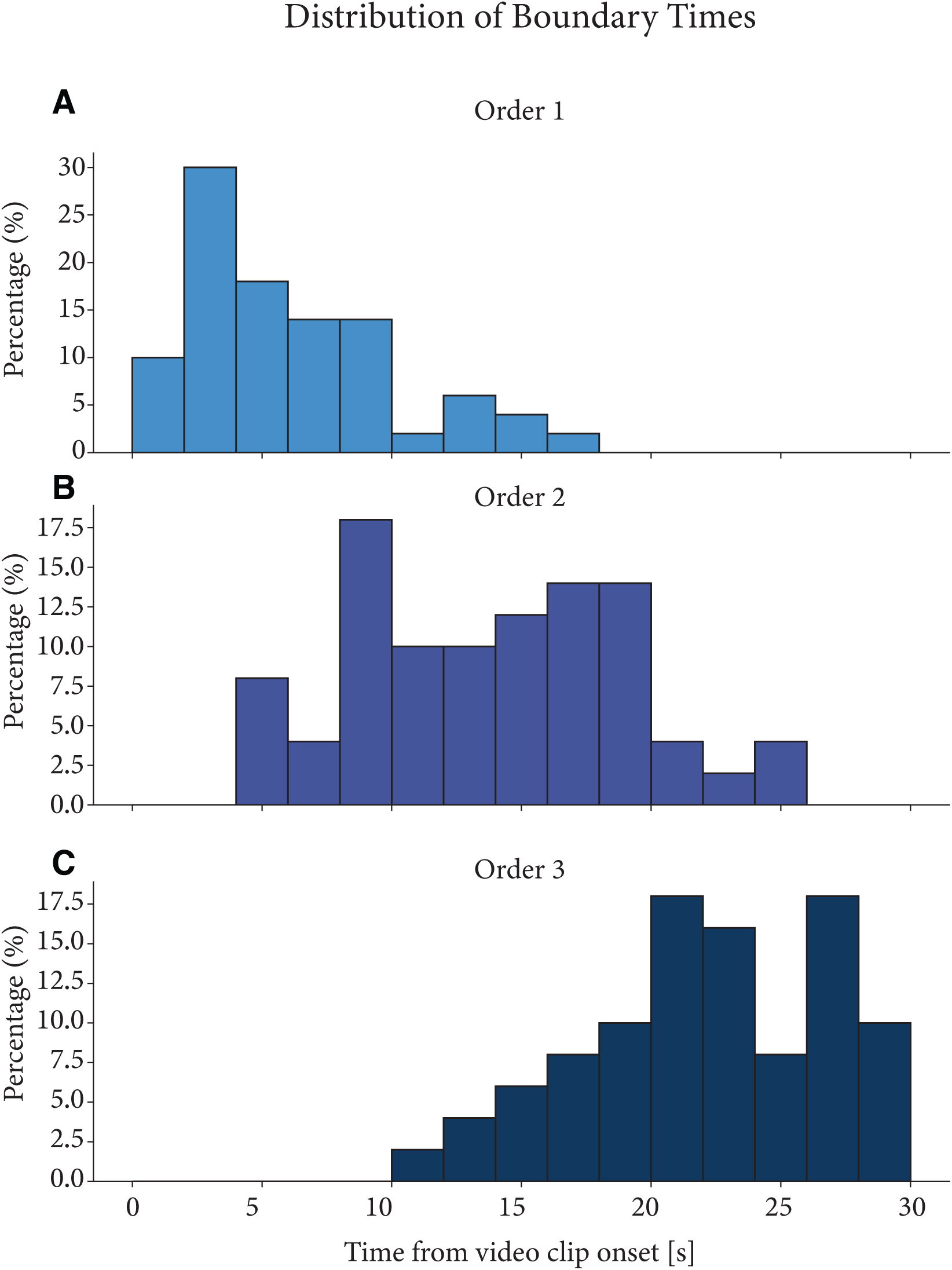
Distribution of event boundary times across video clips. Temporal distribution of the three event boundaries (**A**: Order 1, **B**: Order 2, and **C**: Order 3) pooled across all clips (n = 50 clips total, 25 V1 clips and 25 V2 clips, see *Task* section under *Methods*). Bin size = 2 seconds.

**Figure S2.**
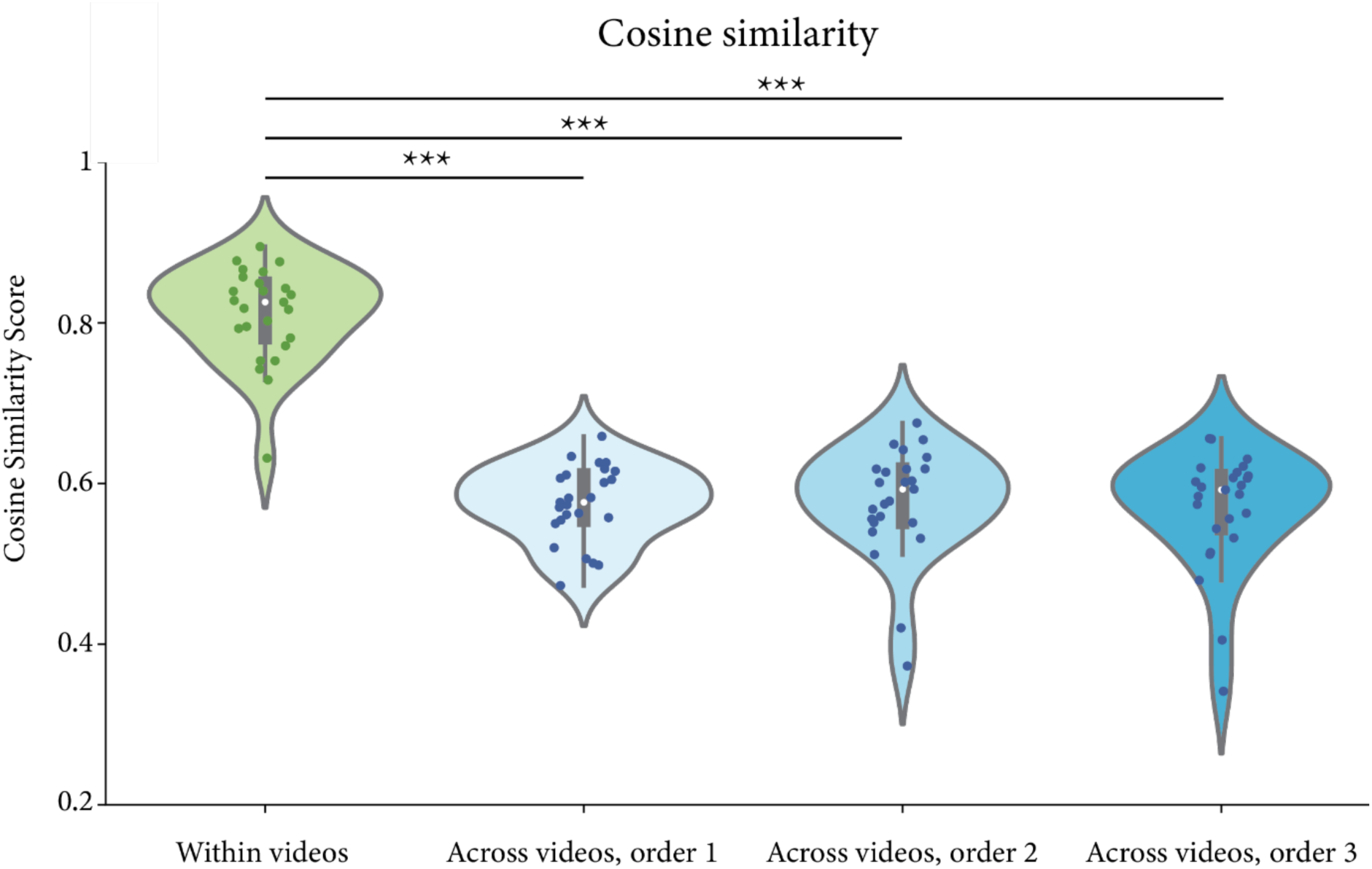
Visual feature similarity was larger within versus across videos. Quantification of visual feature similarity within and across video clips at event boundaries. Feature vectors were extracted from the fc6 layer of AlexNet^18^ for frames at each event boundary. Cosine similarity scores were calculated between frames from the same video (within videos, green) and between frames at corresponding boundaries across different videos (across videos, blue shades). Each dot represents an individual comparison, and asterisks indicate statistical significance when comparing the cosine similarity score between frames within videos than across videos at event boundaries from different ordinal positions (*** *p* < 0.001).

**Figure S3.**
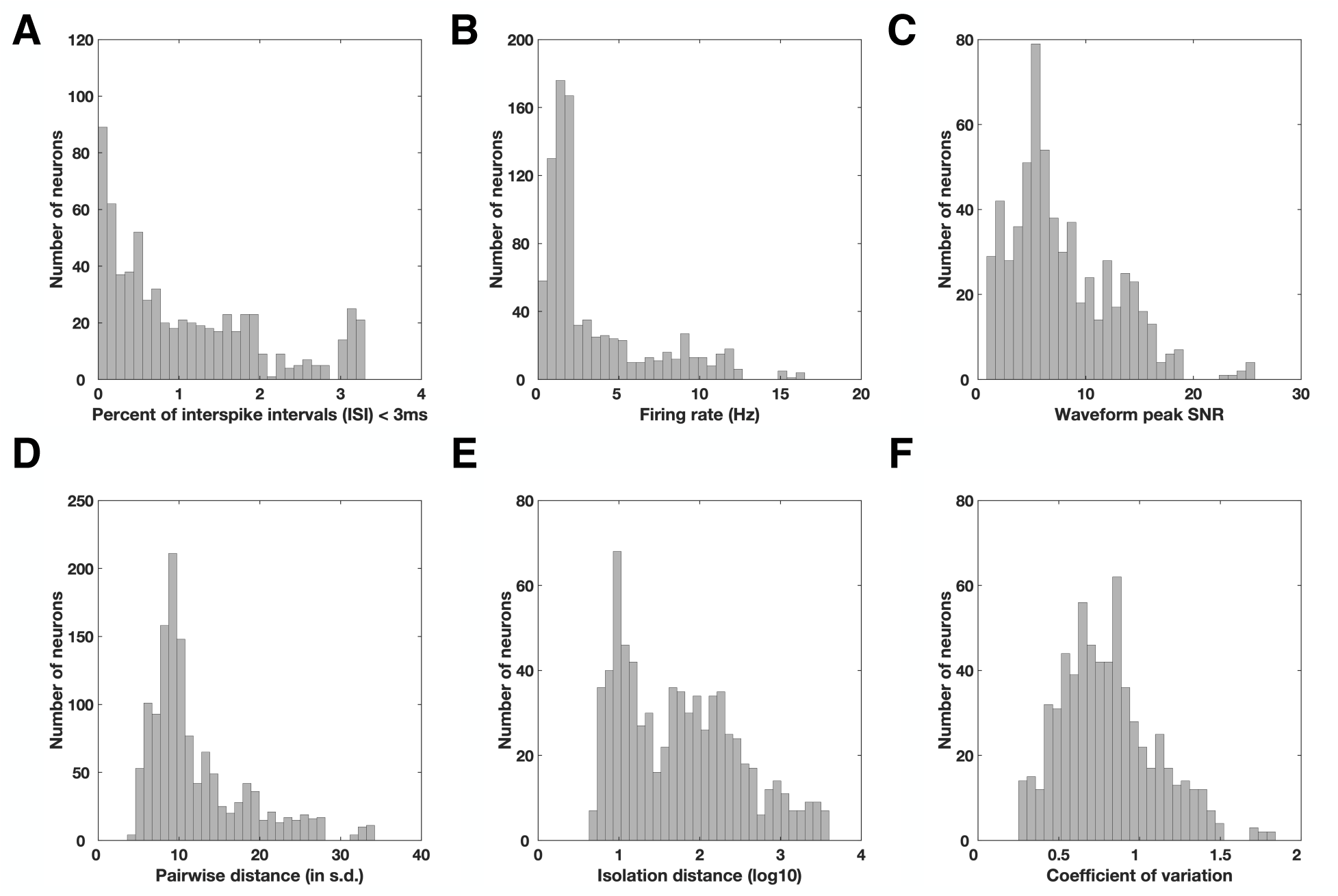
Spike sorting quality metrics. For all identified putative single cells with firing rates higher than 0.5 Hz. **A**, Proportion of inter-spike intervals (ISI) that were shorter than 3ms. **B**, Average firing rate within the entire recording session for all identified putative single cells. **C**, Waveform peak signal-to-noise ratio (SNR), which is the ratio between the peak amplitude of the mean waveform and the s.t.d. of the noise of each identified putative single cell. **D**, Pairwise isolation distance between putative single cells identified from the same wire. **E**, Isolation distance across all identified putative single cells that was calculated in a ten-dimensional feature space of the energy-normalized waveforms. **F**, Coefficient-of-variation (CV2) in the ISI for each identified putative single cell.

**Figure S4.**
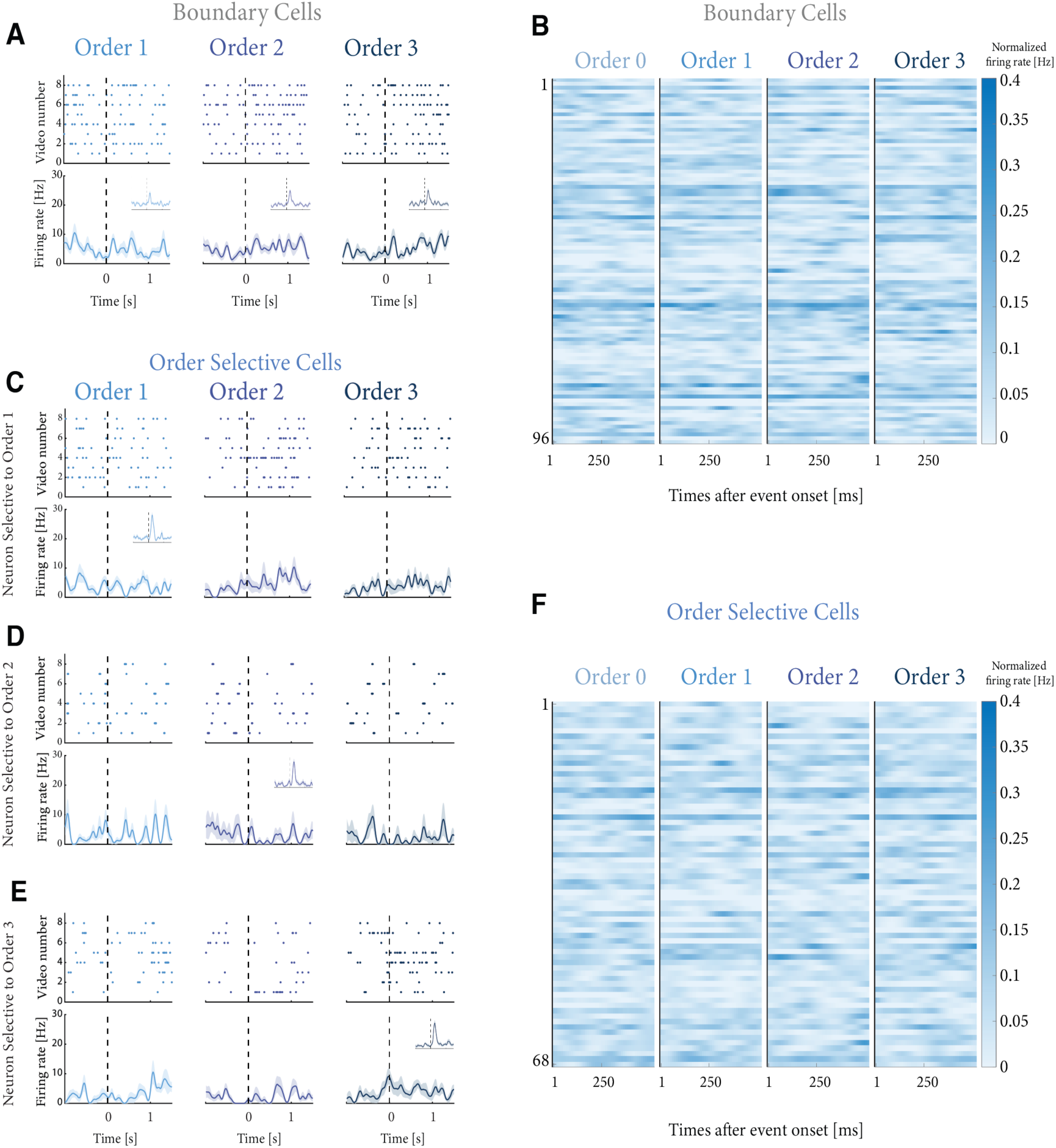
Videos without visual event boundaries do not elicit order selective tuning. **A**, Activity of a boundary cell during trials with no visual cuts. Top: raster plots, bottom: firing rates averaged across a 0.2 s window. Insets show the activity of the same boundary cell during trials with visual cuts. **C, D, E,** Spikes (top row) and mean firing rates (bottom row) of the order selective neuron presented in Figure 2B, now analyzed during control trials in which there are no visual event boundaries. For these trials, the neural activity is aligned to the same moment in time during which an event boundary was introduced in Figure 2B, i.e. at the onset of the event. However, in these control trials, there was no visual boundary introduced at the event boundaries. In the insets, the activity of the same neurons during trials with visual cuts is included. **B** and **F**, Summary plots of the average post-boundary activity of Boundary cells (**B**) and Order Selective Cells (**F**) during trials with no visual event boundaries. Each row is a neuron, and each column is the average activity over 500 ms post-boundary, where the boundary is the onset (column 1), the first, second or third visual boundary (columns 2, 3, 4).

**Figure S5.**
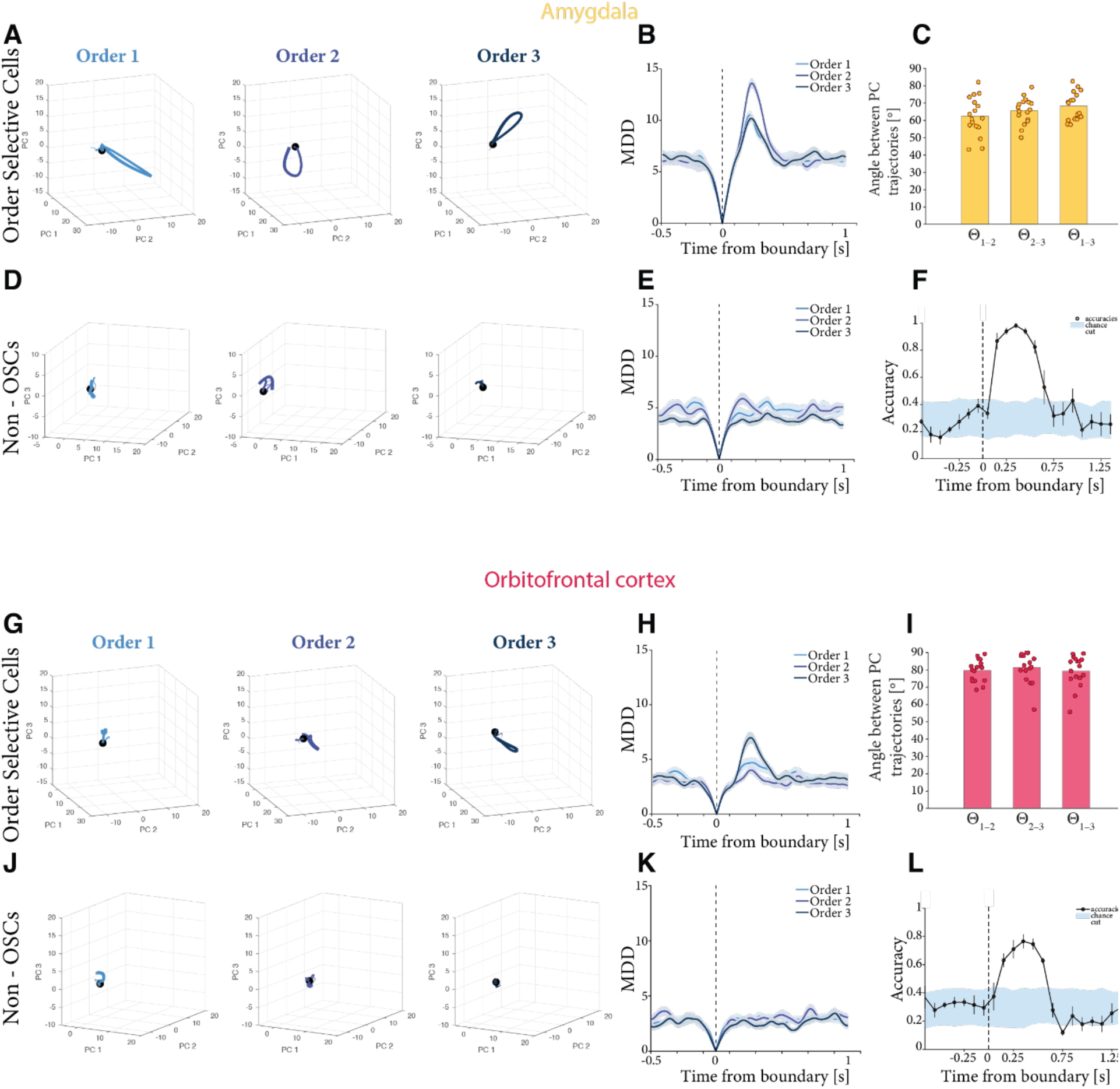
PCA analysis for the amygdala and orbitofrontal pseudopopulations. **A**, Trial-averaged PCA trajectories in time ([-0.5, 0.5] seconds relative to event boundaries) when pooling amygdala neurons classified as OSCs. At the time of the visual cut (black dot), neural states exhibit a large dynamical shift. **B**, Multidimensional Eucledian distance (MDD) between each point of the trajectories in PC space and 0, i.e. when the cut occurred. The shaded area represents ± s.e.m. across trials. **C**, quantification of the angles between the direction of trial trajectories. The bar plot quantifies the average, and the scatter represent the trial-based individual set of angles. **D** and **E**, analogous to A, B, when pooling amygdala neurons that are classified as boundary cells but not order selective. **F**, Decoding performances across time trained on amygdala features. Each dot represents the decoding performance of SVMs trained on 0.5-second windows, with a stride of 0.1 second. The accuracy of each time window is plotted in the middle time point of the time window. In gray is the shuffle variability, while in black are the accuracies. Error bars represent standard errors of the mean cross-validation accuracies. **G**: Trial-averaged PCA trajectories in time ([-0.5, 0.5] seconds relative to event boundaries) when pooling orbitofrontal cortex neurons classified as OSCs. At the time of the visual cut (black dot), neural states exhibit a large dynamical shift. **H**: Multidimensional Eucledian distance (MDD) between each point of the trajectories in PC space and 0, i.e. when the cut occurred. The shaded area represents ± s.e.m. across trials. **I**: quantification of the angles between the direction of trial trajectories. The bar plot quantifies the average, and the scatter represent the trial-based individual set of angles. **J, K**: analogous to G, H when pooling orbitofrontal cortex neurons that are classified as boundary cells but not order selective. **L**: Decoding performances across time trained on orbitofrontal features. Each dot represents the decoding performance of SVMs trained on 0.5-second windows, with a stride of 0.1 seconds. The accuracy of each time window is plotted in the middle time point of the time window. In gray is the shuffle variability, while in black are the accuracies. Error bars represent standard errors of the mean cross-validation accuracies.

**Figure S6.**
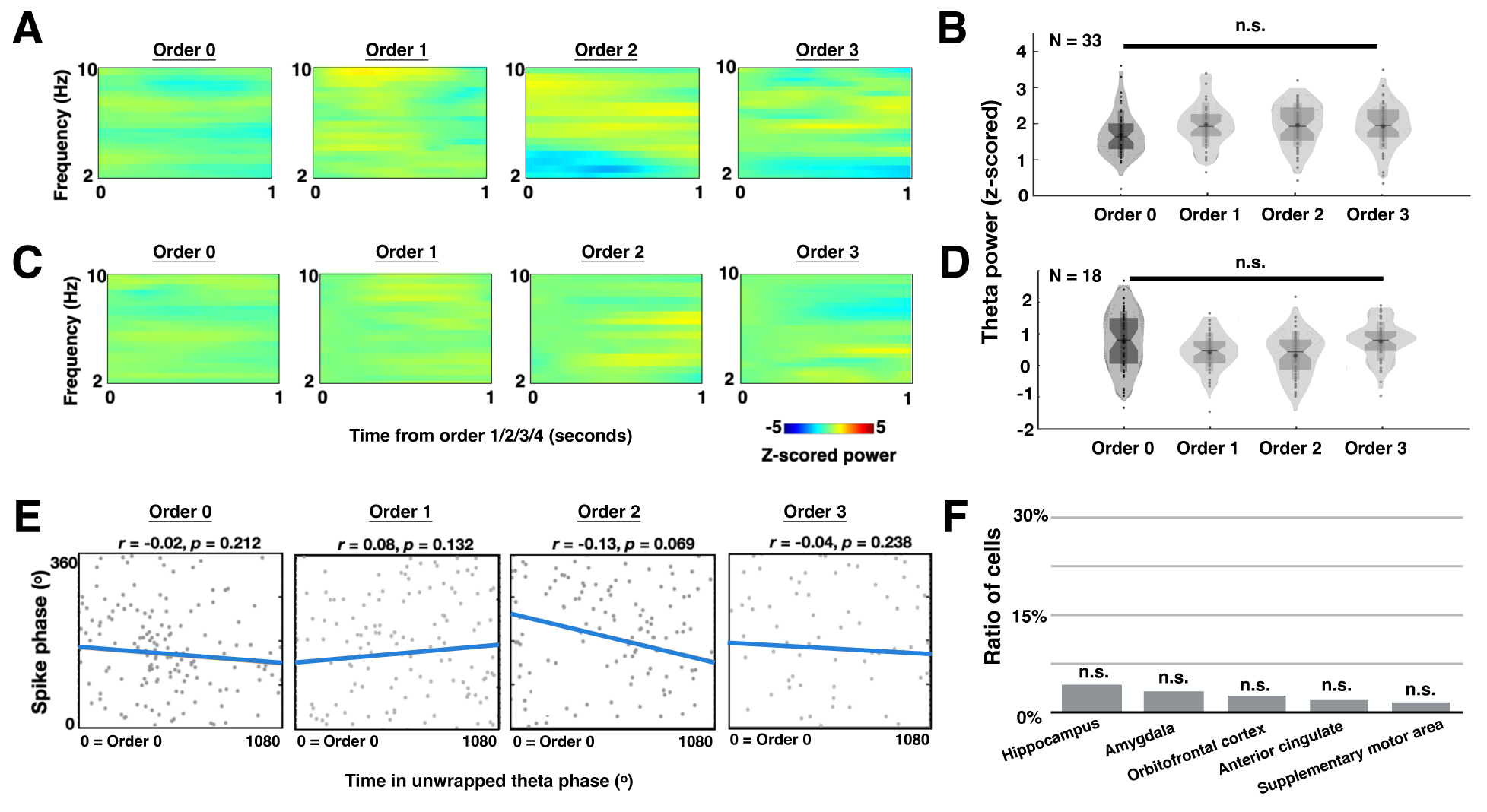
Theta power and phase precession when participants watch no boundary clips. **A, C**, Time frequency plots of the same neuron in Figure 4A and 4C, aligned to the onsets of event boundaries (annotated by the independent group) in the no boundary clips. Power within each frequency band is z-scored and normalized to the baseline period (i.e., fixation cross in Figure 1A). Warmer color denotes power increase relative to the baseline, while colder color indicates power decrease relative to the baseline. **B, D,** Among microelectrodes that demonstrate significant theta power increase (n = 33 electrodes) or decrease (n = 18 electrodes), the distribution of their normalized theta power computed within the 1-second time window following different event boundaries in the no boundary clips. *n.s. = not significant*, ANOVA test, see Methods. **E,** Example order selective neuron in Figure 4E shows no theta phase precession following the onset of different event boundaries in no boundary clips. The strength of theta phase precession is quantified as the correlation coefficient (*r*) between time in unwrapped theta phase and spiking phases relative to the underlying theta oscillations. Note that more negative correlation coefficients denote stronger theta phase precession. **F,** Number of neurons showing significant phase precession during encoding when participants watch no boundary clips. *n.s. = not significant*, permutation test, see Methods.

**Figure S7.**
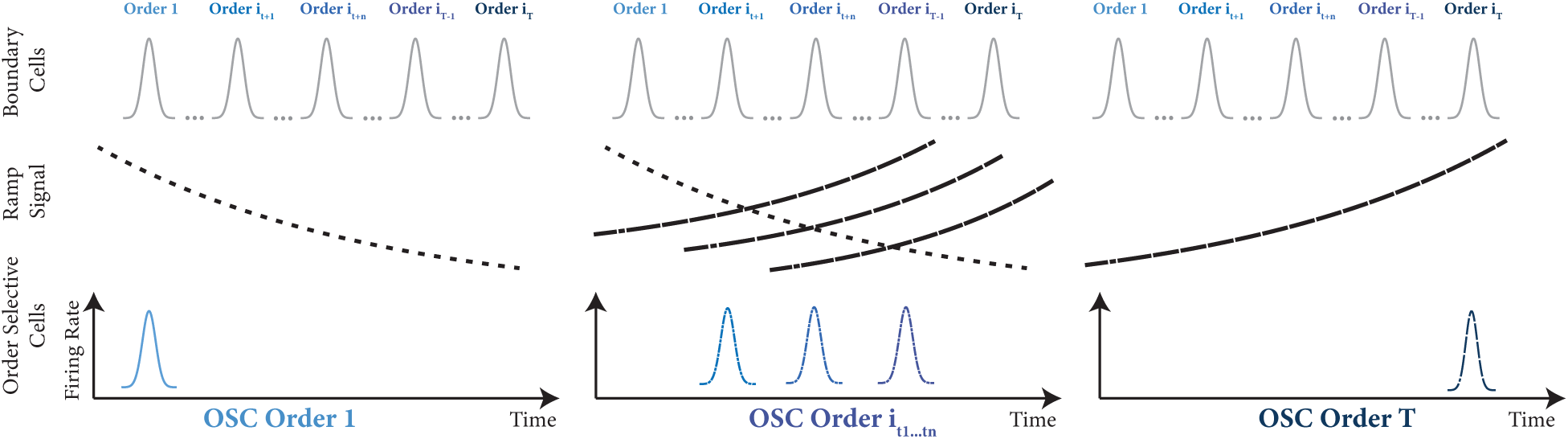
Schematic conceptual model illustrating the proposed mechanistic origin of OSCs. The top row represents boundary cells that respond to all event boundaries regardless of order. The middle row depicts time-dependent signals within each trial segment, reported in other work^12^. The bottom row shows how order-selective responses emerge from the product between boundary detection and temporal signals, where the different orders may arise depending on the nature of the time-varying ramp. This framework could explain with first principles how neurons develop selective responses to boundaries at specific orders within event sequences while maintaining invariance to visual content.

**Table S1.** Patient information and distribution of OSCs and Boundary cells. Note that P79CS_v1 and P79CS_v2 are data collected from the same patients across different days.

| Patient ID | Subject ID | Age | Gender | All cells (965) | OSC cells in HPC (27) | OSC cells in AMY (25) | OSC cells in OFC (8) | OSC cells in other regions (8) | Boundary cells (96) |
| --- | --- | --- | --- | --- | --- | --- | --- | --- | --- |
| P78CS | 1 | 54 | F | 23 | 0 | 0 | 0 | 0 | 4 |
| P79CS_v1 | 2 | 49 | F | 109 | 0 | 3 | 2 | 2 | 11 |
| P79CS_v2 | 2 | 49 | F | 54 | 0 | 2 | 0 | 0 | 5 |
| P80CS | 3 | 24 | O | 41 | 0 | 0 | 0 | 0 | 1 |
| P81CS | 4 | 28 | F | 48 | 1 | 1 | 1 | 0 | 8 |
| P82CS | 5 | 42 | M | 32 | 2 | 1 | 0 | 0 | 6 |
| P87CS | 6 | 26 | F | 56 | 1 | 3 | 0 | 1 | 8 |
| P88CS | 7 | 29 | F | 125 | 3 | 2 | 3 | 1 | 8 |
| AMC20 | 8 | 60 | F | 21 | 0 | 0 | 0 | 0 | 4 |
| AMC21 | 9 | 22 | F | 14 | 0 | 0 | 0 | 0 | 1 |
| AMC23 | 10 | 34 | F | 9 | 0 | 0 | 0 | 0 | 0 |
| AMC24 | 11 | 27 | F | 8 | 0 | 3 | 0 | 0 | 2 |
| TWH202 | 12 | 27 | M | 10 | 0 | 0 | 0 | 0 | 0 |
| TWH206 | 13 | 23 | M | 85 | 5 | 3 | 0 | 1 | 15 |
| TWH207 | 14 | 45 | F | 46 | 3 | 2 | 0 | 1 | 4 |
| TWH208 | 15 | 33 | M | 58 | 5 | 0 | 0 | 0 | 5 |
| P90CS | 16 | 32 | M | 26 | 0 | 0 | 0 | 0 | 0 |
| P92CS | 17 | 30 | F | 83 | 3 | 5 | 0 | 0 | 7 |
| TWH209 | 18 | 35 | M | 35 | 0 | 0 | 0 | 1 | 1 |
| TWH212 | 19 | 63 | M | 22 | 0 | 0 | 0 | 0 | 1 |
| TWH214 | 20 | 40 | M | 60 | 4 | 0 | 2 | 1 | 5 |

**Table S2.** MNI coordinates of microwire bundles on which at least one OSC was detected.

| Electrode ID | Patient ID | Location | Side | MNI Coordinates<br>X | MNI Coordinates2<br>Y | MNI Coordinates3<br>Z |
| --- | --- | --- | --- | --- | --- | --- |
| 52, 55, 56 | P79 v1 | amygdala | right | 25.3 | 0.78 | -32.34 |
| 194, 196 | P79 v1 | orbitofrontal cortex | left | -1.04 | 41.25 | -16.91 |
| 24 | P79 v2 | amygdala | left | -16.85 | 1.02 | -26.28 |
| 55 | P79 v2 | amygdala | right | 25.3 | 0.78 | -32.34 |
| 23 | P81CS | amygdala | left | -19.98 | -2.51 | -23.18 |
| 27 | P81CS | hippocampus | left | -17.28 | -14.76 | -15.24 |
| 202 | P81CS | orbitofrontal cortex | right | 1.63 | 33.13 | -23.09 |
| 179 | P82CS | amygdala | right | 16.7 | -4.1 | -16.98 |
| 186, 192 | P82CS | hippocampus | right | 25.95 | -16.91 | -13.97 |
| 145 | P87CS | amygdala | left | -22.76 | -13.09 | -22.6 |
| 160 | P87CS | hippocampus | left | -26.59 | -26.17 | -21.03 |
| 162 | P87CS | anterior cingulate | right | 7.42 | 32.3 | 20.78 |
| 178, 184 | P87CS | amygdala | right | 20.81 | -7.19 | -22.7 |
| 146 | P88CS | amygdala | left | -18.33 | -5 | -17.31 |
| 153, 160 | P88CS | hippocampus | left | -26.37 | -16.92 | -15.9 |
| 162 | P88CS | anterior cingulate | right | 10.84 | 29.14 | 29.08 |
| 185 | P88CS | hippocampus | right | 25.08 | -18.03 | -16.95 |
| 193, 195, 198 | P88CS | orbitofrontal cortex | left | -2.17 | 36.17 | -11.15 |
| 3, 4 | AMC24 | amygdala | left | -15.61 | -7.33 | -16.58 |
| 6 | TWH206 | hippocampus | left | -28.03 | -29.83 | -12.39 |
| 44, 45 | TWH206 | hippocampus | left | -24.27 | -38.52 | -5.07 |
| 85, 86, 88 | TWH206 | amygdala | right | 25.06 | -4.96 | -20.84 |
| 120 | TWH206 | hippocampus | right | 25.21 | -38.62 | -4.33 |
| 92 | TWH207 | insula | left | -36.78 | 12.59 | -20.89 |
| 102 | TWH207 | amygdala | left | -21.99 | -5.05 | -28.3 |
| 107, 111 | TWH207 | hippocampus | left | -26.84 | -39.08 | -3.25 |
| 141 | TWH207 | amygdala | right | 27.43 | -4.1 | -22.6 |
| 66, 68 | TWH208 | hippocampus | left | -23.16 | -19.79 | -21.31 |
| 87 | TWH208 | hippocampus | left | -22.03 | -43.18 | -8.74 |
| 98, 100 | TWH208 | hippocampus | right | 25.39 | -16.52 | -18.09 |
| 148, 150 | P92CS | amygdala | left | -16.28 | -3.39 | -22.11 |
| 157 | P92CS | hippocampus | left | -22.1 | -12.78 | -21.29 |
| 180, 184 | P92CS | amygdala | right | 14.07 | -4.29 | -23.17 |
| 185, 190 | P92CS | hippocampus | right | 18.48 | -12.22 | -22.82 |
| 156 | TWH214 | orbitofrontal cortex | left | -4.03 | 9.33 | -22.02 |
| 162, 167, 168 | TWH214 | hippocampus | left | -26.87 | -23.57 | -11.8 |
| 186 | TWH214 | hippocampus | right | 27.28 | -14.94 | -15.49 |
| 213 | TWH214 | orbitofrontal cortex | right | 3.25 | 8.89 | -24.2 |

